# 3D projection electrophoresis for single-cell immunoblotting

**DOI:** 10.1101/805770

**Authors:** Samantha M. Grist, Andoni P. Mourdoukoutas, Amy E. Herr

## Abstract

While immunoassays and mass spectrometry are powerful single-cell protein analysis tools, bottlenecks remain in interfacing and throughput. Here, we introduce highly parallel, synchronous, three-dimensional single-cell immunoblots to detect both cytosolic and nuclear proteins. The novel threedimensional microfluidic device is a photoactive polyacrylamide gel with a high-density microwell array patterned on one face (x-y) for cell isolation and lysis. From each microwell, single-cell lysate is ‘electrophoretically projected’ into the 3^rd^ dimension (z-axis), separated by size, and photo-captured for immunoprobing and three-dimensional interrogation by confocal/light sheet microscopy. Design guidelines for throughput and separation performance are informed by simulation, analyses, and deconvolution postprocessing based on physics of 3D diffusion. Importantly, separations are nearly synchronous, whereas serial analyses can impart hours of delay between the first and last cell. We achieve an electrophoresis throughput of >2.5 cells/s (70X faster than serial sampling) and perform 25 immunoblots/mm^2^ device area (>10X increase over previous immunoblots). A straightforward device for parallel single-cell immunoblotting, projection electrophoresis promises to advance integration of protein-level profiles into the emerging single-cell atlas of genomic and transcriptomic profiles.

## Introduction

Proteins are biomolecules that play a direct role in nearly all cellular processes^1^. As such, protein expression is a primary metric for quantifying cell state^2^. Although genomics and transcriptomics analysis of gene and RNA expression are powerful and complementary measurement techniques, they often do not predict protein expression at the single-cell level^3–5^. Proteoforms, or different forms of proteins arising from the same gene^6^, are also critical to understanding cell state: cellular heterogeneity at the proteoform level plays critical roles in cellular processes including regulating tumor growth^7^ and resistance to treatment in cancer^8^. Single-cell proteomic technologies have emerged for characterizing protein expression heterogeneity at the single-cell level, but many such technologies are limited to protein detection by antibodies with limited protein specificity, which may not be proteoform-specific^9^. Protein separations based on electrophoresis can overcome these antibody specificity limitations through dual measurements of the physicochemical properties of protein targets (i.e., molecular mass, isoelectric point) and primary antibody reactivity for protein detection.

To offer a comprehensive understanding of complex cellular systems, selective detection of protein targets in individual cells is required; yet, key single-cell analysis challenges persist in throughput, sample preparation, and interfacing^10^. Measurement throughput is crucial to detect rare but important subpopulations of cells (e.g., metastasis, resistance to treatment), but – to attain suitable statistical power – the minimum sample size increases as subpopulation prevalence decreases^11^. Thus, the ability to assay hundreds or even thousands of cells is valuable. Further compounding the challenge of measuring a large cell population with single-cell resolution is synchronously assaying all of the single cells. Proteomic cell state is dynamic^12^ so differences in analysis time across cells can add artefactual heterogeneity – particularly when assessing cell response to drugs or environmental stimuli. Furthermore, single-cell lysate preparation and interfacing for interrogation must also preserve protein concentrations from dilution: the total protein copy number in a single cell can be low due to the small cellular volume, even for the most highly expressed proteins. Diffusion into larger-than-a-cell volumes, sample losses, and lysate changes during transfer to an analysis platform can further challenge single-cell proteoform measurement^10^.

Since the early 1990’s, capillary electrophoresis (CE) has worked to surmount challenges in analysis of lysate from individual cells^13^. To maintain protein target concentrations during analysis, CE uses glass capillaries with micron-scale diameters, facilitating efficient dissipation of Joule heating and ultra-rapid electrophoresis. For lysate transfer, the CE systems use physical alignment of the capillary to reagent baths or isolated single cells. Early single-cell CE technologies were capable of assaying 10’s of cells per day, making measurement of single-cell heterogeneity among large cell populations out of reach^14,10^. Improvements in CE throughput use capillary arrays for simultaneous analyses of samples. For example, Capillary Array Electrophoresis (CAE) uses 48 glass capillaries run in parallel with two separations per capillary to achieve 96 separations in 8 min^15^. While parallelization boosts throughput, CAE device fabrication is intensive, involving design and alignment of electrode, reservoir, and capillary arrays. Bundled microstructured silica fibers support up to 168 simultaneous separations, but from one bulk sample (demonstrated to improve heat transfer rather than throughput)^16^. A challenge to the use of glass CE technologies for single-cell measurement is that they can suffer from analyte loss via adsorption to the capillary surface due to large surface-area-to-volume ratios^10^.

To advance single-cell resolution analyses, researchers brought both automation and microfluidic design to bear to efficiently integrate disparate sample preparation and electrophoretic analysis. In an excellent example, the Allbritton group introduced an automated, serial single-cell electrophoresis system by integrating a single glass capillary with a microwell array for cell isolation, using a motorized stage for microwell alignment to the capillary for serial sampling and analysis^17^. For analysis of a population of 219 mammalian cells, throughput was clocked at 2.1 cells/min (cell lysis, electrophoresis, and real-time fluorescence detection of sphingosine fluorescein and sphingosine-1-phosphate fluorescein). Also exemplar of integrated approaches, planar microfluidic devices support analysis of up to 12 cells/min, with notable reductions in reagent consumption, as compared to conventional systems^18,19^. While useful, serial analysis of large cell populations is inherently asynchronous and, as mentioned, can introduce artefactual cell-to-cell heterogeneity, particularly critical to assessing a response to drug or another stimulus.

Mass spectrometry is a powerful, complementary separations technology that is promising for single-cell protein detection. Aside from mass cytometry^20^, other forms of mass spectrometry do not rely on antibody specificity for selective protein detection. Highly multiplexed protein detection from single cells (1000’s of targets per cell) is possible with bottom-up mass spectrometry, although throughput has not yet exceeded 100 individual cells^21,22^. Mass spectrometry is limited to highly abundant proteins (>10^4^ copies per cell)^21,22^ and small-to-intermediate molecular masses for top-down proteomics (e.g. typically 25 kDa for MALDI-TOF, although specialized detectors facilitate detection of proteins up to 110 kDa^23,24^). Furthermore, the vast majority of single-cell mass spectrometry approaches are bottom-up, wherein the requirement for protein digestion to peptides confounds proteoform stoichiometry. While top-down mass spectrometry does measure intact proteins – making the approaches relevant to proteoforms – and imaging mass spectrometry can assay larger numbers of cells (e.g. >1000 dorsal root ganglia)^25^, larger proteins present a measurement challenge even with optimized sequential analysis for wider-range mass measurement (m/z 400 to 20 000)^25^.

Immunoblotting is a class of separations with a powerful capacity for targeted proteomics. To detect protein targets *a priori* identified through hypotheses or discovery tools, this suite of separations approaches integrate two analytical modalities to yield enhanced target specificity over either alone: separation of proteins by electrophoresis and probing of specific targets by immunoreagents. While only recently developed for single-cell and sub-cellular resolution by our group, immunoblotting (and its most popular form, western blotting) have been a workhorse in biological and clinical laboratories for decades. Using automation in a different way to advance towards both single-cell resolution and multiplexed detection, the Kennedy group designed an automated system that integrates microchip electrophoresis with immunoprobing on an off-chip PVDF membrane^26^. Integration between the microchip and membrane uses a mobile membrane, with the separations effluent deposited (blotted) onto that moving blotting membrane. While not demonstrated for single-cell analysis, multiplexing was boosted to 11 separated protein targets from 9 serial separations in 8 min, with each separation from the same aliquot of bulk cell lysate with a total protein content similar to that of a single cell (400 ng total protein). Taking a different approach inspired by single-cell DNA electrophoresis (COMET assays)^27^, we introduced single-cell western blotting with a throughput of ~200 cells/min^28^. The planar 2D device uses a thin layer of polyacrylamide gel stippled with microwells for parallel cell isolation and lysis. Protein lysate is subjected to immunoblotting in the photoactive polyacrylamide gel abutting each microwell. We have applied the single-cell western blotting technology to studying heterogeneity of circulating tumour cells^29^, smooth muscle cells^30^, and HER2 isoform expression in clinical specimens^31^. Furthermore, using similar design principles we have introduced single-cell immunoblotting tools to separate proteins based upon other physicochemical protein properties (e.g. subcellular localization^32^ and isoelectric point^33,34^) and adapted the single-cell western blotting assay to assess adherent cells attached to the gel surface (without detachment)^35^ and study invasive motility of glioblastoma cells^36^. These tools have emerged as powerful technologies for single-cell analysis; however, their throughput and sample consumption remain limited by the large spacing between microwells required for the protein separation^35^.

Here, we introduce a novel high-density parallelized single-cell immunoblotting device that uses the third dimension (z-axis), to enhance the microwell array density and, hence, throughput and sample consumption. In projection electrophoresis, we sought to leverage the full gel volume to map protein size and originating cell position from the 3D location of separated protein bands. The 3D design is inspired by multifunctional 3D hydrogel materials like Expansion Microscopy^37^ and CLARITY^38^, as well as the efficient electrotransfer of protein from gel to PVDF membrane in conventional western blotting^39,40^ and bulk separations in layered systems^41,42^. Using a 3D volume improves microwell array density to 25 microwells/mm^2^ from 2 microwells/mm^2^ in 2D (planar) devices. Owing to a small microwell pitch in projection electrophoresis, the assay consumes an order of magnitude less volume of a cell suspension (25 μL vs. 300 μL) to assay ~300 cells (similar to planar systems), resulting in a 10-fold improvement in sample consumption. In this work, we describe the separation performance and use the physics of 3D diffusion to design the projection electrophoresis system, and perform near-simultaneous immunoblotting of both cytoplasmic (GAPDH, actinin, beta tubulin) and nuclear (PTBP1) protein targets from hundreds of single mammalian breast and brain tumour cells.

## Results and Discussion

### Establishing projection electrophoresis as an analytical tool

In lieu of serial interrogation and electrophoretic analysis of individual mammalian cells, projection electrophoresis (**Figure 1**a) yields synchronous, concurrent analyses of hundreds of single cells. Compared to serial cell measurements, the parallel approach reduces assay-induced protein expression heterogeneity (**Figure 1**b). After in-gel immunoprobing for the protein targets glyceraldehyde 3-phosphate dehydrogenase (GAPDH, involved in glycolysis, transcription, and apoptosis) and polypyrimidine tract-binding protein 1 (PTBP1, RNA-binding protein involved in cellular processes including splicing), projection electrophoresis yields 3D data (with X-Y position describing the originating microwell, Z-position describing protein size, **Figure 1**c-d). Mean Z-direction migration distances were 199±9 μm for PTBP1 (57 kDa) and 346±7 μm for GAPDH (37 kDa). Furthermore, the multifunctional, micropatterned polyacrylamide gel (PAG) serves as the cell isolation device, the lysis vessel, the separation matrix, and the protein capture scaffold. This multifunctional gel thus facilitates *in situ* lysis, electrophoresis, and blotting, mitigating losses incurred in sample transfer steps, while synchronous analysis and fast assay times yield high assay throughput and drastically reduced sampling delays between cells in a population **Figure 1**e).

**Figure 1.**
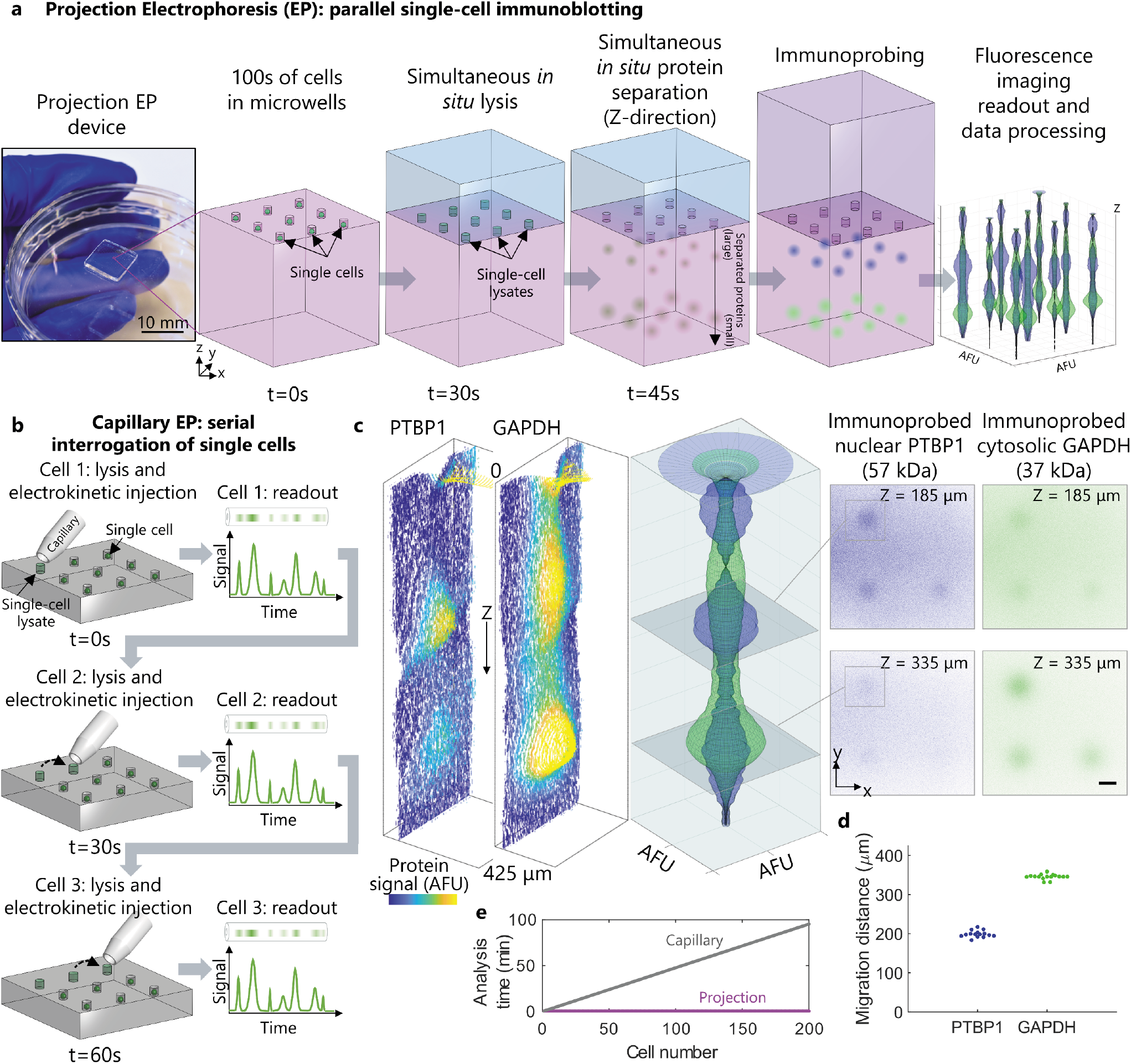
Projection electrophoresis simultaneously lyses and separates both nuclear and cytosolic proteins from hundreds of single cells. **(a)** Projection electrophoresis device photograph and workflow schematic. A 9×9 mm projection electrophoresis polyacrylamide gel contains >1000 microwells that each serve as a separation lane. Cells in polyacrylamide gel microwells are lysed *in situ* using a lysis buffer-soaked hydrogel delivery matrix, and lysate is then electrophoretically injected into the polyacrylamide and proteins separate by size through the depth of the gel. After separation, proteins are covalently linked to the gel matrix using a UV-initiated capture. All cells are lysed and analyzed simultaneously, the active assay time (from cell lysis to photocapture) is less than 90s, and *in situ* analysis reduces potential losses from transfer steps. **(b)** In contrast, capillary electrophoresis analyzes cells in series, with each cell lysed and proteins separated at a different time. **(c)** Visualization of Z-directional protein separation in a single separation lane. (left) X-Z (Y-summed) contour plots of background-subtracted protein signal within the separation lane for nuclear and cytosolic example proteins. Protein signal is visualized as peak height and false-coloured. (middle) Z-directional intensity profiles (summed fluorescence intensity vs. Z) for PTBP1 (nuclear, blue) and GAPDH (cytosolic, green) in the same lane, revolved around the Z-axis to generate a 3D rendering of fluorescence distribution. (right) Confocal X-Y slice images for separated PTBP1 and GAPDH at two Z-depths into the gel (top: through the PTBP1 band; bottom: through the GAPDH band). Each slice image shows 4 separation lanes, 3 of which were occupied by BT474 cells prior to analysis. Overlaid squares depict X-Y regions of interest for the lane plotted in the left and middle sub-panels, over which fluorescence intensities were summed to yield Z-directional intensity profiles. Scale bar represents 50 μm. **(d)** Quantified Z-migration distances for PTBP1 (57 kDa) and GAPDH (37 kDa) from n=15 (PTBP1) and n=17 (GAPDH) separation lanes. **(e)** Comparison of cell-cell variation in cell lysis/analysis time for serial capillary electrophoresis (throughput of 2.1 cells/min) and projection electrophoresis. While cells are lysed at times varying by over 1.5 hours with serial interrogation, all cells are lysed and proteins separated within 1.5 minutes in projection electrophoresis.

We first sought to verify the separation mechanism governing protein electrophoresis in the concurrent analyses (**Figure 2**a-b). To understand the protein separation mechanism, we assessed electromigration of a ladder of well-characterized protein standards (donkey immunoglobulin anti-mouse IgG, IgG: 150 kDa; bovine serum albumin, BSA: 66.5 kDa; ovalbumin, OVA: 42.7 kDa, each labelled with AlexaFluor^®^ dyes). When purified protein solution is pipetted on top of the gel block (with gel block face stippled with microwells), the protein solution preferentially partitions into the microwells (versus the hydrogel), thus providing a convenient, well-controlled means for sample loading into each microwell sample injector. To minimize 3D diffusional spreading during PAGE, we designed an ultra-short separation axis (1-mm) and rapid (<1 min) protein PAGE duration. The length of the PAGE separation axis is defined by the gel block thickness. Upon completion of PAGE, the multifunctional gel was toggled from separation matrix to a protein capture scaffold using a 45-s exposure to UV illumination^43^.

**Figure 2.**
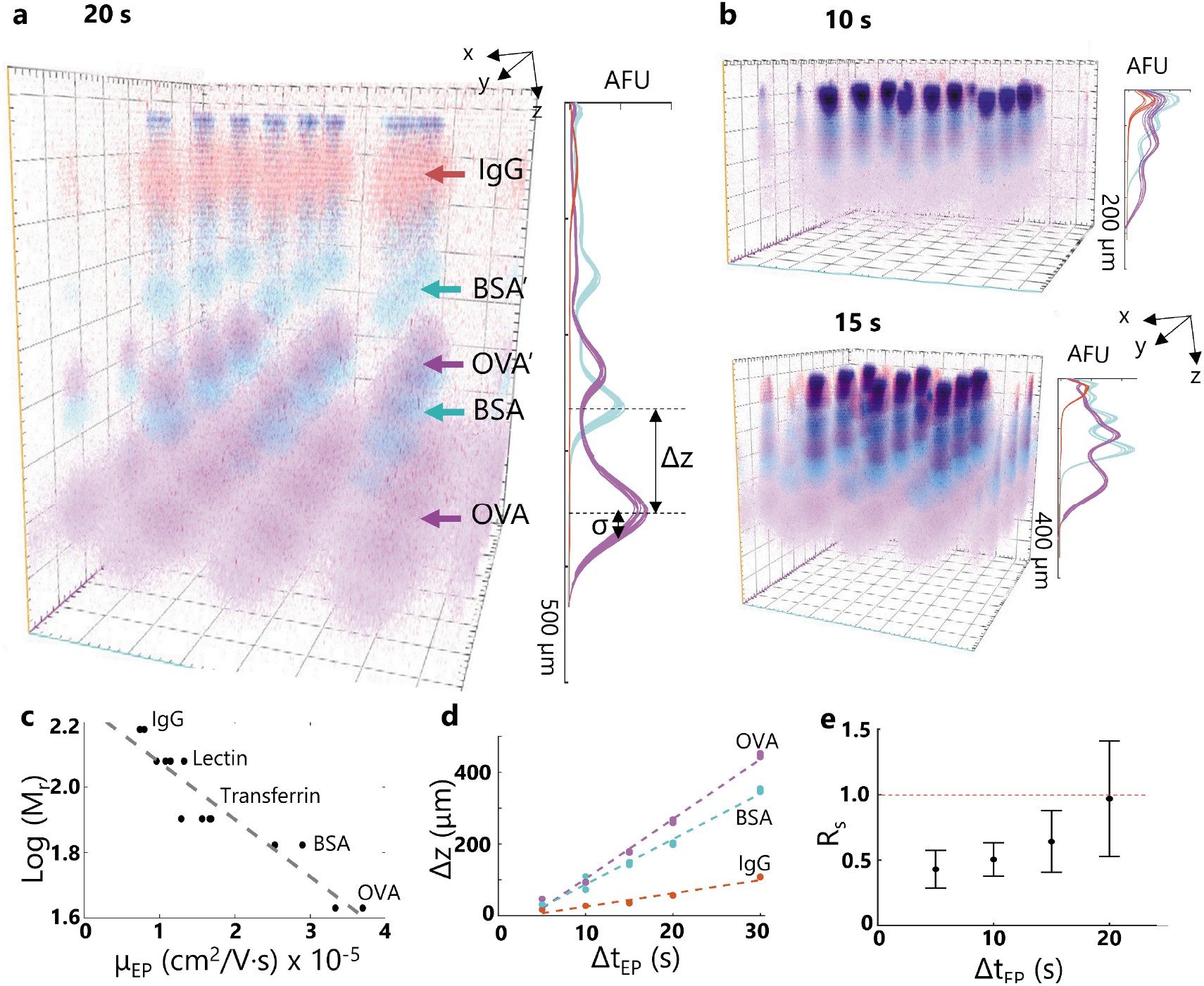
Projection electrophoresis supports protein PAGE. **(a)** Confocal imaging of PAGE of fluorescently labeled protein ladder at 20 s elapsed separation time. Each ladder sample is injected from a 32 μm diameter microwell (x-y plane) with PAGE along the z-axis of the gel block. Summing x-y separation lane intensities (10%T PA separation gel) yields z-intensity profiles, with peak-to-peak displacement Δz and peak width σ. **(b)** 3D renderings and Z-intensity profiles are plotted for (i) 10s electrophoresis and (ii) 15s electrophoresis in 10%T polyacrylamide gels for comparison with the data for 20s electrophoresis shown in (a). **(c)** The electrophoretic mobility of the proteins depends log-linearly on protein molecular weight. Each plotted point represents an electrophoretic mobility calculated from linearly fitting migration distance vs. electrophoresis time data from 11-17 segmented separation lanes in n=5 gels (5 electrophoresis times). Linear fitting yields log(M_r_)=1.7×10^4^μ_EP_+2.25 (R^2^=0.89). **(d)** Electromigration distance depends linearly on electrophoresis time, thus proteins migrate at constant velocity during PAGE. Migration distances are plotted from protein bands originating from 11-17 nearby microwells in two independent 7%T polyacrylamide gels containing 10% Rhinohide; 33 mA constant current; 52 V/cm initial, 2 gels for each migration time. Linear fitting yields OVA migration=16.7t-63.42; BSA migration=12.64t-39.3; IgG migration=3.69t-11.54. **(e)** Separation resolution R_s_ for BSA and OVA peaks in 10%T PAGE gels. Each point depicts the mean and standard deviation of the R_s_ calculation from the median migration distances and peak widths from n=4 independent separation gels.

We observed (**Figure 2**c) a log-linear relationship of electromigration with molecular mass, as expected in size-sieving gel electrophoresis^44^. Further, in this unique format, we observed constant-velocity and size-dependent electromigration for the ladder and additional protein assayed (**Figure 2**d; electrophoretic mobilities of OVA: 3.3-3.7×10^-5^ cm^2^/Vs; BSA: 2.5-2.9×10^-5^ cm^2^/Vs; transferrin: 1.3-1.7×10^-5^ cm^2^/Vs, lectin: 0.96-1.14×10^-5^ cm^2^/Vs; IgG: 7.4-8.0×10^-6^ cm^2^/Vs). Constant-velocity migration required mitigation of deleterious effects of electrolysis (i.e., buffer pH changes, bubble formation at the electrodes), which increased the R^2^ of linear fits to the protein migration data from 0.87±0.06 to 0.97±0.03 for the three ladder protein species (Figure S1, Supplementary Information). For BSA and OVA ladder species, both a protein monomer and dimer are resolvable, as would be expected in high performance protein PAGE^45–48^. Having established the separation mechanism, we next estimated the PAGE performance by assessing the separation resolution (R_S_). For two ladder proteins (OVA, BSA) in a 1-mm thick 10%T PA gel volume (**Figure 2**e), the Rs exceeded 1.0 within 20 s of PAGE, yielding fully resolved species.

Based on the dominant separation mechanism and rapid protein separation, analysis of the purified protein ladder solution suggests that projection electrophoresis is suitable for analytical-quality protein analysis. The high performance of the rapid microfluidic protein analysis described here is in contrast to another 3D system, designed for preparatory Z-direction separation performance, as previously demonstrated for bulk samples using a multilayered gel to coarsely fractionate small proteins (14-77 kDa) from large proteins (20-343 kDa)^42^. In terms of throughput, each purified protein projection electrophoresis gel (100 μm microwell pitch) contains >4000 microwells, facilitating >4000 parallel (replicate) purified protein separations for a total active assay throughput of 44 separations per second (not including readout time). For comparison, Capillary Array Electrophoresis of a single sample yields a throughput of 5 separations per second^15^.

### Device design and imaging approach are informed by 3D diffusion of target proteins

Given the open microfluidic design of the projection electrophoresis device that uses a microwell array to perform sample isolation and preparation, with an abutting gel volume that performs the analytical functions (protein PAGE, immunoblotting), we sought to understand physics-based factors that set the minimum acceptable microwell-to-microwell spacing (microwell pitch, Δ_well_). The microwell pitch, in turn, sets the maximum number of concurrent protein PAGE separations per projection electrophoresis device. As illustrated schematically in **Figure 3**a, the Δ_well_ spacing is influenced by the length scale of diffusional band broadening (σ_xy_, in X-Y plane) during protein PAGE along the Z-axis. As design guidelines, the throughput of each single-cell projection electrophoresis device (number of single cells assayed per device) will be a function of Δ_well_ (sets separation lane density) and overall usable device dimensions (**Figure 3**b). As Δ_well_ depends linearly on σ_xy_, the maximum lane density is inversely proportional to σ_xy_^2^, as computed in **Figure 3**c. Two design rules are plotted: at Δ_well_ > 4σ_xy_, we would estimate <5% protein overlap between neighboring lanes, while at the more conservative Δ_well_ > 6σ_xy_, we would estimate <0.3% protein overlap assuming Gaussian protein distributions.

**Figure 3.**
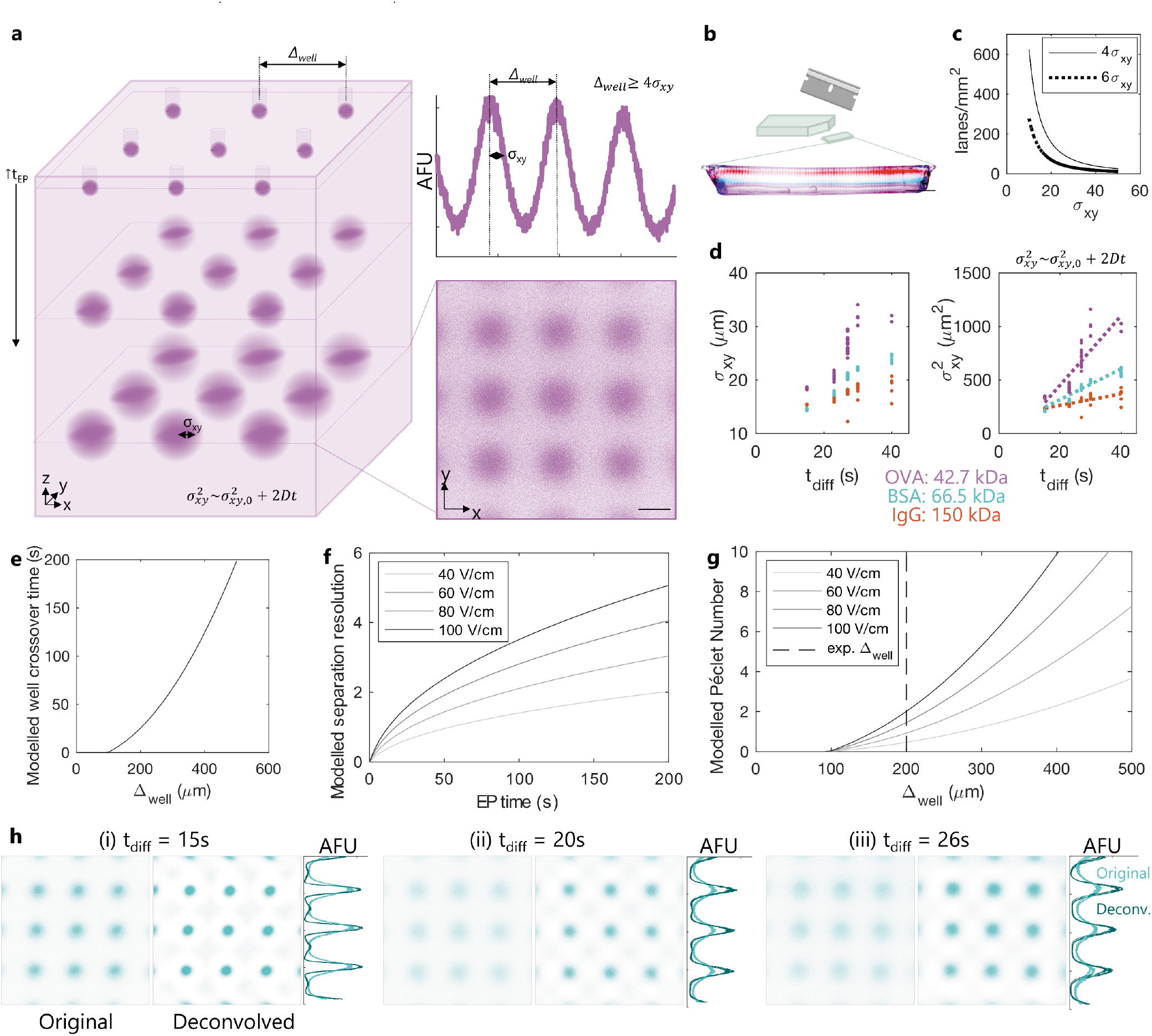
The physics of 3D diffusion dictate Projection EP device design and inform image analyses. **(a)** Lane density (limited by minimum spacing between separation lanes Δ_well_) is dependent on separated protein x-y band width (σ_xy_) to avoid microwell-microwell crosstalk. σ_xy_ in turn depends on 3D diffusion of the injected protein. Simulated data shown in left schematic; measured OVA data shown in micrograph and intensity profile. Scale bar represents 100 μm. **(b)** Throughput is a function of both lane density and usable gel area; separated protein bands parallel to the gel edges in a cross-sectional image of the gel show uniform migration across the gel. Scale bar represents 1 mm. **(c)** Theoretical maximum lane density at a spacing of 4σ_xy_ (~5% overlap between lanes) and 6σ_xy_ (~0.3% overlap between lanes. Maximum lane density is inversely proportional to σ_xy_^2^. **(d)** Measured diffusional x-y band broadening (Gaussian fit peak width σ_xy_) from purified proteins initially partitioned into 32 μm microwells, as a function of in-gel diffusion time (t_diff_). Left: σ_xy_ vs. t_diff_. Right: σ_xy_^2^ vs. t_diff_ with linear fits (*σ_xy_^2^ = ρ_xy,0_^2^+2D*t_diff_). After 10s electrophoresis, we measure σ_xy_ < 30 μm for all proteins, suggesting that 200 μm microwell spacing will be sufficient. Linear fitting yields OVA σ_xy_^2^ =32.5t_diff_-198 (R^2^=0.75); BSA σ_xy_^2^ =14.3t_diff_+29.8 (R^2^=0.88); IgG σ_xy_^2^ =5.41t_diff_+153 (R^2^=0.42). **(e)** Modelled time for ovalbumin bands to cross over into the neighboring lane, as a function of microwell spacing Δ_well_. **(f)** Modelled separation resolution for the model protein pair BSA and OVA, as a function of electrophoresis time and electric field strength. **(g)** Modelled Péclet number (defined as the ratio of the time to reach a separation resolution of 1 for the model protein pair BSA and OVA to the time at which the protein band may be expected to cross over into the neighboring separation lane. **(h)** Physics-driven postprocessing. For each confocal slice example (BSA, 7%T gel), the original image, that after physics-driven postprocessing (deconvolution of a point spread function modeling 3D diffusion), and summed intensity profiles for a 100 μm region surrounding each row of protein bands in both the original and deconvolved images are shown. Each image pair is scaled to the maximum of the (higher-intensity) deconvolved image.

During electromigration, protein peaks will diffuse in three dimensions, with diffusion along the Z-axis determining separation resolution (R_S_) and diffusion in X-Y determining the minimum Δ_well_. Diffusional spreading of protein bands in all three dimensions depends on protein molecular mass, temperature, time, and the gel density (pore size)^49,50^. To assess the impact of protein diffusion on setting Δ_well_, we assessed the well-characterized fluorescently labeled OVA/BSA/IgG protein ladder during protein PAGE in a 7%T gel projection electrophoresis device. For each time point analyzed by confocal imaging, we determined the Z position of the maximum of the summed fluorescence intensity, for an X-Y region of interest surrounding the each microwell injector. At this Z position, we assessed X-Y resolution by Gaussian fitting X and Y intensity profiles and extracting the mean fitted peak width σ_xy_. For duplicate gels of 5 electrophoresis times, we plotted the squared peak width vs. time in gel, fitting to the expected diffusional peak spreading^49^:

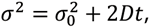

For each protein target, the *σ_0_* is related to the injected peak width (dictated by microwell diameter), *D* is the in-gel diffusion coefficient, and *t* is the total elapsed time since protein injection. **Figure 3**a shows an example confocal fluorescence data X-Y slice image for OVA and associated Δ_well_ design rule. Applying the analysis to the full protein ladder (**Figure 3**d), yields estimates of Δ_well_, across a range of protein targets and diffusion coefficients (D_OVA_ ~16 μm^2^/s, D_BSA_ ~ 7 μm^2^/s, D_IgG_ ~ 2.7 μm^2^/s calculated from linear fits to the plot of σ_xy_^2^ vs. diffusion time). Under the described conditions, the protein target with the largest *D* (OVA) suggests that a Δ_well_ of 200 μm will satisfy the trade-off of maximizing separation lane density while minimizing separation lane overlap (7%T gels, 30 s protein PAGE). For comparison, top-down MALDI imaging mass spectrometry utilizes a protein spot pitch of 20-200 μm^24^.

To further understand the diffusion-driven interdependency of the microwell spacing and separation performance, for the highest diffusivity ladder protein (OVA) we modelled the maximum assay time (the time at which protein signal is expected to bleed into the neighboring lane, from the diffusional peak spreading function above) for a range of microwell spacings (**Figure 3**e), as well as the *R_S_* as a function of electrophoresis time and electric field strength (**Figure 3**f). The separation resolution *R_S_* is modeled as^51^:

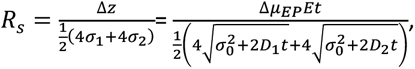

where *Δz* is the difference in migration distance between the two protein targets, *σ_1_* and *σ_2_* are the Z-direction peak widths, *Δ_μEP_* is the difference in electrophoretic mobility between the two targets, and *D_1_* and *D_2_* are the diffusion coefficients for the two targets. Diffusion coefficients are provided in Table 1 of the Methods, with electrophoretic mobilities empirically determined in **Figure 2**d.

**Table 1.** Protein hydrodynamic radii (*r_H_*), diffusion coefficients (*D*) and partition coefficients (*k*) used in finite-element modelling of diffusion during lysis and electrophoresis.

| Protein | $r_H$[nm] | $D_{\text{sol}}$<br>[m<sup>2</sup>/s] | $D_{7\%T}$<br>[m<sup>2</sup>/s] | $D_{20\%T}$[m<sup>2</sup>/s] | $k_{7\%T}$ | $k_{20\%T}$ |
| --- | --- | --- | --- | --- | --- | --- |
| turboGFP | 2.3 | $6.61 \times 10^{-11}$ | $1.36 \times 10^{-11}$ | $9.49 \times 10^{-13}$ | 0.501 | 0.0415 |
| BSA | 3.01 | $5.05 \times 10^{-11}$ | $7.90 \times 10^{-12}$ | $3.49 \times 10^{-13}$ | 0.344 | 0.00744 |
| HER2 | 3.76 | $4.04 \times 10^{-11}$ | $4.88 \times 10^{-12}$ | $1.39 \times 10^{-13}$ | 0.213 | $8.16 \times 10^{-4}$ |

We next defined a Péclet number as the ratio of the maximum assay time to the time required to reach *R_S_*=1, with results presented in **Figure 3**g and the Péclet number given by:

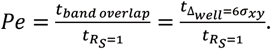

At Δ_well_ = 200 μm (**Figure 3**d), Pe ~1 for an applied electric field strength of 60 V/cm. As protein band diffusion measurements (**Figure 3**d) suggest measured diffusion coefficients smaller than predicted values, our Péclet analysis is a conservative estimate of the trade-off between separation performance and achievable microwell density. Increasing the electric field strength increases the separation resolution and, thus, the Péclet number. While we demonstrate electric fields of 43-68 V/cm in this work, high electric fields are achievable with reasonable applied voltages in this system due to the small (0.3 cm) electrode spacing (100 V/cm only requires V_app_ = 30 V, compared to 100s to 1000s of V for typical electrophoresis configurations).

Building on understanding of the dominant physics, namely diffusion, we next sought to investigate computational approaches to recovery of starting concentration distributions (in the microwell array) from endpoint confocal fluorescence images of the protein PAGE (**Figure 3**h). In microscopy, deconvolution of an experimentally, theoretically, or computationally-determined, microscope-dependent point spread function (psf) from 2D or 3D images recovers spatial resolution by image postprocessing^52–54^. Inspired by deconvolution in microscopy, we explored whether we could represent the final protein projection image (*I(x,y,z,t)*) as the initial protein x-y pattern (*p(x,y,z)*; the spatial arrangement of cells/microwells) convolved with a ‘diffusional point spread function’ *psf_diff_(x,y,z,t)*, in turn convolved with the imaging point spread function *psf_img_(x,y,z,t)*:

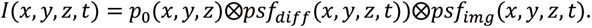

We chose to describe the diffusional *psf* using 3D point-source diffusion^55^:

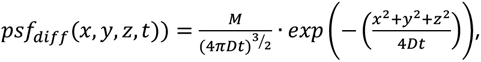

where *D* is again the protein diffusion coefficient, *t* the elapsed diffusion time, and *M* the starting number of molecules at the point source. Although we estimated the full 3D point spread function for each protein, we performed 2D deconvolution only on the individual slice images, without using information from neighboring focal planes (as in a “no-neighbors” deconvolution imaging method^56^). We used this approach to simplify processing, while recognizing that the simplification limits a full 3D reconstruction and the signal intensity improvement possible from 3D deconvolution. To perform 2D processing, we deconvolved the 2D function *psf_diff_* (through the centre of the point spread function, at Z=0) from 2D confocal slice images at the Z-direction migration peak for each protein (the Z position at which the summed intensity for that protein in the image field of view was maximized). We neglected effects of the imaging point spread function, as we expect that the resolution of our measurement is more limited by diffusion (10s of microns for our typical time scales, as shown in **Figure 3**a) than by the resolution of confocal microscopy with high NA objectives (typically sub-micron^57^).

After deconvolution of the protein PAGE images, we observe a considerable improvement in spatial resolution of XY profiles of separated BSA (**Figure 3**h; 5-15s elapsed PAGE duration in (i)-(iii)). Comparing the ‘original’ to the ‘deconvolved’ images illustrates that spatial resolution is reduced from σ_xy_ = 16±2 μm to σ_xy_ = 8.4±0.2 μm (47%) for 5s electrophoresis (15s total time until photocapture), σ_xy_ = 25.7±0.6 μm to σ_xy_ = 13.1±0.6 μm (49%) for 10s electrophoresis (15s total time), and σ_xy_ = 26.3±0.6 μm to σ_xy_ = 13.3±0.5 (49%) for 15s electrophoresis (26s total time). Further, the localization of the peak centre was unperturbed by reconstruction (Δμ < 1.1 μm for all analyzed protein spots, with Δμ_avg_ = 0.38 μm) and the integrated fluorescence signal of each protein sample is minimally perturbed by the reconstruction except when visible artefacts were present in the deconvolved images as shown in the lowest electrophoresis time (i) (average AUCs after postprocessing are within 2.5% of the initial values in (ii-iii), but 24% in (i)). Measured errors in peak center were Δμ=0.08±0.06 μm (t_diff_=15s), Δμ=0.68±0.3 μm (t_diff_=20s), Δμ=0.1±0.2 μm (t_diff_=26s); measured errors in peak AUCs were ΔAUC=23.9±1.4% (t_diff_=15s), ΔAUC=0.5±0.8% (t_diff_=20s), ΔAUC=3±2% (t_diff_=26s) (n=9 ROIs). Taken together, we see physics-based image postprocessing as a promising approach to reconstructing a map of a starting sample from the target concentration distributions in the 3D gel volume, using the endpoint fluorescence readout of protein PAGE.

### Design for integration of single-cell sample preparation into projection electrophoresis

Having considered design of the projection electrophoresis device and assay using a well-characterized protein ladder, we next sought to identify factors important to high-performance protein PAGE of single cells (**Figure 4**). We first assessed settling of single BT474 cells in 25 μm diameter microwells within the projection electrophoresis gels. **Figure 4**a depicts settled Calcein-stained BT474 breast tumour cells in microwells, alongside a corresponding full-gel wide-field microscopy image of immunoprobed GAPDH fluorescence after the projection electrophoresis assay; probed protein bands correlate with settled cell positions. Cell settling efficiencies (populated microwells) were at 43±8% with the number of settled single cells 356±82 per 9×9 mm projection electrophoresis device (n=5 devices). The fraction of microwells occupied by more than one cells was 10±3%. Further optimization of cell settling densities, microwell geometries, and settling times would likely improve these values and thus assay throughput.

**Figure 4.**
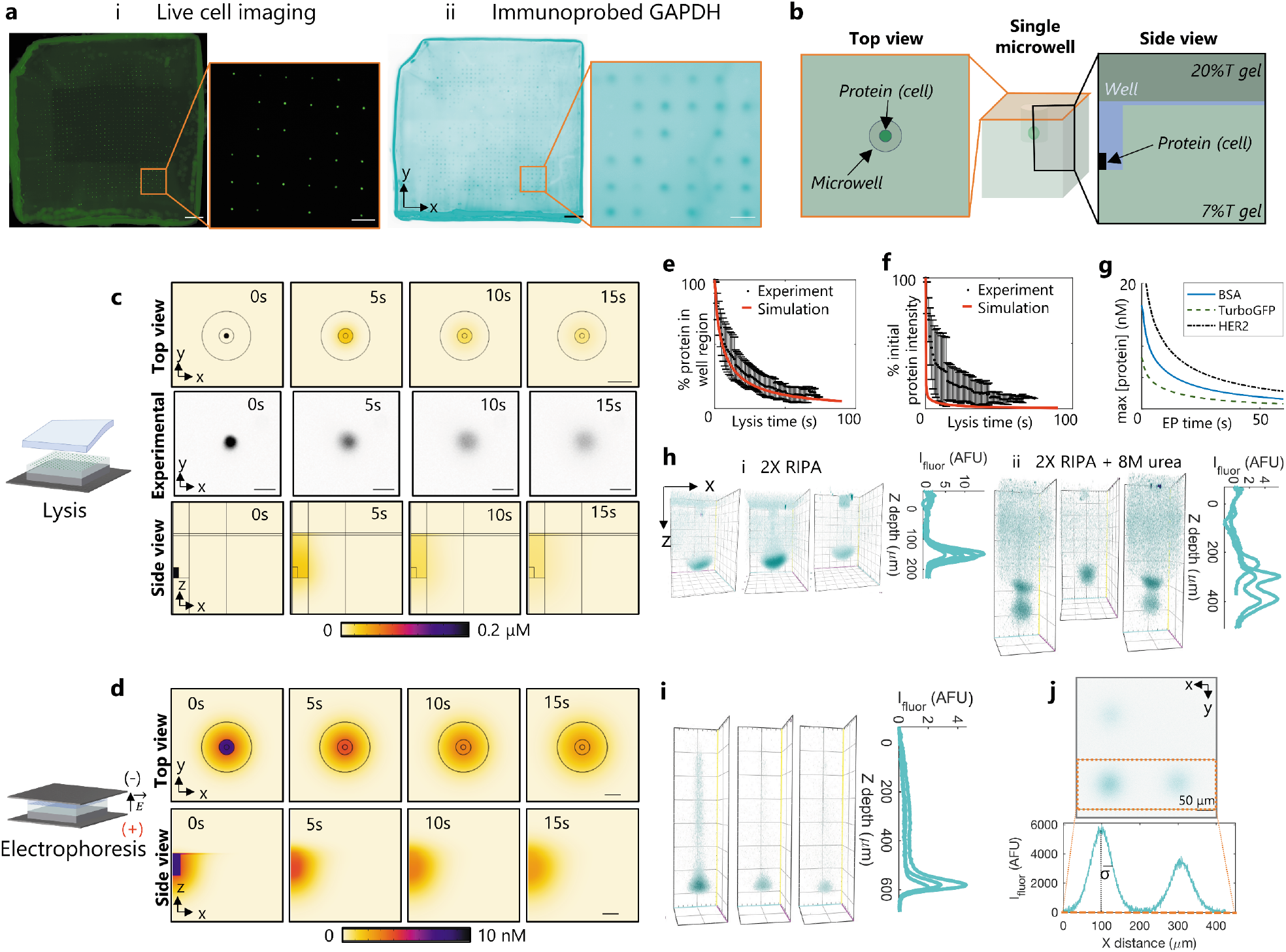
Design and verification of sample preparation for projection electrophoresis of single mammalian cells. (**a**) High-density endogenous protein bands (ii) correspond to single-cell settling in microwells (i). Scale bars represent 1 mm (left full-gel images) and 200 μm (right zoom images). (**b**) Illustration of top-view and side-view geometries shown in protein dilution studies (c-d). (**c**) Modelling and experimental quantification of diffusional dilution during lysis. Simulated and experimental top-view images of diffusional protein dilution during lysis, and side-view simulated results are shown. Simulated initial TurboGFP concentration was 2 μM. Scale bars represent 50 μm. (**d**) Modelling the impact of diffusion during electrophoresis on detectable in-gel protein concentration. Side and top view TurboGFP concentration profiles are shown at different times during electrophoresis. Simulated initial TurboGFP concentration (before lysis and electrophoresis) was 2 μM. Scale bars represent 30 μm. **(e)** Quantification of the percent of protein remaining in the microwell region during lysis. **(f)** Quantification of the change in the spatial maximum protein concentration as a function of time after protein solubilization. **(g)** Simulated maximum protein concentration vs. electrophoresis time, for 3 model proteins. (**h**) Representative beta tubulin separations from U251 glioblastoma cells lysed with different buffer formulations, both after 10s electrophoresis. Lysis/EP buffer requires 8M urea for fast protein solubilization and electromigration. **(i)** Maximum intensity projection 3D renderings and Z-intensity profiles of probed GAPDH bands from single BT474 breast tumour cells. **(j)** Microwell packing density (impacting assay throughput) is dependent on protein band diffusion before photocapture. Protein diffusion profiles confirm that a microwell pitch of 200 μm is sufficient to resolve bands from neighboring microwells. After 10s EP, the mean peak width (σ) of the X-Y GAPDH spots is 32±13 μm. At a microwell pitch of 192 μm (3σ), <0.3% of the signal should bleed into the neighboring lane. Scale bar represents 50 μm.

After protein solubilization, diffusion-driven dilution of single-cell lysate occurs rapidly in the open microwell geometries. To determine how the concentration of the single-cell protein lysate changes during the lysis and electrophoresis stages of the assay, we used a combination of finite-element modelling (geometries shown in **Figure 4**b) and experimental monitoring of turboGFP-expressing U251 cells during cell lysis (**Figure 4**c) and finite-element modelling during electrophoresis (**Figure 4**d). To compare with wide-field microscopy imaging data of turboGFP cell lysis, we integrated the 3D protein concentrations over the full Z range of the model to mimic detected wide-field fluorescence intensities (**Figure 4**e-f). Using this metric, after a typical 25s lysis time, we measure 17%±11% of the initial protein intensity. Our simulated profiles overestimate the diffusional dilution of protein, predicting only 2.2% of the initial intensity for in-well lysis after 25s. We attribute the ~15% discrepancy to a possible diffusion barrier on the microwell wall surface, arising from either the presence of Rhinohide^®^ in the gel matrix, or the presence of residual GelSlick^®^ or dichlorodimethylsilane used during gel fabrication. One important consideration for projection electrophoresis is that, in contrast to other single-cell protein separation assays such as scWB^20,28,31,32,58–60^ and scIEF^34,61,62^, there is reduced protein ‘loss’ in in the Z direction projection platform –primarily dilution. While other electrophoretic cytometry assays have a fluid layer or lid gel above the thin separation gel, into which protein can diffuse and is lost, in the projection electrophoresis device most (if not all) protein is mobilized into the bulk of the 3D gel when an electric field is applied to initiate PAGE.

We then sought to quantify how the maximum protein concentration changes during the analytical single-cell PAGE stage (**Figure 4**g). Using finite-element modelling, we first modelled 25s of lysis before using the resulting concentration profile as the initial condition for a model of in-gel diffusion. From an initial protein concentration of 2 μM in a cylinder representing the cell, we estimate maximum protein concentrations of 2.1 nM (turboGFP, 26 kDa), 4.0 nM (BSA, 66.5 kDa), and 6.8 nM (HER2, 185 kDa) after 25s lysis and 20s electrophoresis in **Figure 4**g. We compared the expected diffusional dilution during electrophoresis to that expected in a planar system (**Figure S2**, Supplementary Information) and found similar dilution during the assay steps in both systems. We note that the planar system is amenable to imaging during electrophoresis, thus our comparison (which predicts similar losses to those reported in our previous work^51^) serves as validation of the numerical model. From simulations of X-Y spot size from 3D diffusion of β-tubulin, we expect that the protein concentration 80 μm away from the band centre will be <5% of the maximum concentration at the peak of the protein spot, confirming that 200 μm microwell spacing should also be sufficient for endogenous proteins from single cells. Diffusional dilution of protein is dependent on both analyte size and gel density. The relatively large-pore-size gels used in this work (7%T) are optimal for large analytes (80-200 kDa), with adaptation for smaller analytes accommodated by moving to higher-density (smaller pore size) separation gels.

In optimizing the projection assay, we sought buffer chemistries to minimize lysis and solubilization times and used diffusive immunoprobing of model proteins β-tubulin and GAPDH using immunoglobulin fragments (F(ab) fragments) to assess solubilization efficacy. Here, we assessed a range of cell lysis and protein solubilization chemistries (**Figure 4**h). Across a range of chemistries, we observed differences in protein electromigration and dispersion, which were dependent on buffer composition and delivery methods. We selected a dual-function lysis and solubilization buffer that utilizes the anionic detergents sodium dodecyl sulphate (SDS) and sodium deoxycholate, augmented with a strong chaotrope (8M urea). Comparing solubilization, electromigration, and dispersion of the model protein β-tubulin from U251 glioblastoma cells lysed both without (i) and with (ii) 8M urea in the lysis buffer, we observed rapid electromigration into the 3D gel from the microwell and lower protein peak dispersion with urea present. Without urea, the 3D protein bands exhibited a hollow, bowl-like shape (concave towards the microwell), rather than the 3D Gaussian distribution which would be expected from diffusion theory. In a sub-set of separation lanes, two β-tubulin peaks were detectable after solubilization with urea lysis buffer (**Figure 4**h), suggesting delayed solubilization for a subset of the β-tubulin molecules, as might be expected depending on the intracellular state of the β-tubulin.

In formulating design guidelines for the dual-function lysis/solubilization and electrophoresis buffer, we consider two additional points. First, detergents such as SDS and Triton X-100 form micelles of size on the order of nanometers^63,64^. Consequently, we explored the corollary hypothesis that size-exclusion partitioning^65,66^ of solutes from polyacrylamide gels may limit delivery of such molecules into the bulk of the gel. As the density of polyacrylamide gel negatively correlates with in-gel concentration of size-excluded species^65^, we explored whether lower density (6%T vs. 20%T) polyacrylamide lysis gels may facilitate improved protein solubilization. By moving to 6%T lysis gels, we observed higher apparent GAPDH mobility (1.08±0.03×10^-4^ cm^2^/V·s using 6%T lysis gel, compared with 0.83±0.08×10^-4^ cm^2^/V·s using 20%T lysis gel, n=12-14 separation lanes) and potential reduction in dispersion of the protein band towards the microwell (representative separation lane profiles with each lysis gel condition are in **Figure S3**). Second, strong chaotropes like urea solubilize proteins by disrupting hydrogen bonds as well as electrostatic and hydrophobic interactions to unfold hydrophobic protein regions^67^. Urea-based lysis buffers can solubilize different subsets of the proteome, as compared to RIPA-like buffers^68^. High concentrations of urea (e.g., 8 M) can break down detergent micelles and disturb detergent-protein complexes^69,70^. Urea, as a small molecule, is less susceptible to size-exclusion partitioning from hydrogels. Just as in other protein separations, the ideal lysis buffer depends on the system and target of interest^67^. Analysis of another endogenous target protein, GAPDH, after lysis and protein PAGE in in the 8M urea lysis buffer showed protein peaks with low dispersion (**Figure 4**i).

Lastly, we verified the device design suggested by analysis of the well-characterized protein ladder now for the analysis of mammalian cells using the optimized cell preparation protocol (**Figure 4**j). We anticipated that the cell preparation steps and required time may increase lysate dilution away from that assessed using the idealized protein ladder system in **Figure 3**, both because diffusion of protein targets in each single-cell lysate occurs during the time required for lysis and solubilization and because single-cell PAGE is run at higher temperature (37°C vs. 4°C) to improve protein solubilization. As discussed above, microwell spacing dictates the achievable sample multiplexing on one device. After 10s EP, we measured σ = 32±13 μm for GAPDH in the X-Y plane. At a Δ_well_ = 192 μm (6σ), we estimate that <0.3% of the fluorescent signal from the GAPDH in each cell lysate should bleed into the neighboring lane. Using the device and sample preparation protocols designed here, we observed protein diffusion profiles for endogenous GAPDH that confirm that selection of Δ_well_ = 200 μm is sufficient to limit cross-contamination between adjacent separation lanes.

### Projection electrophoresis for immunoblotting of protein targets from hundreds of single mammalian cells

We applied projection electrophoresis to immunoblotting analyses of well-characterized endogenous proteins GAPDH and actinin across populations of individual human BT474 breast cancer cells. As depicted in **Figure 5**, projection electrophoresis concurrently analyzes hundreds of single cells by parallelized separation after near-simultaneous lysis. While in the initial characterization of **Figure 4** we used labelled F(ab) fragments rather than full-length antibodies to mitigate long diffusive timescales and size-exclusion partitioning that challenge in-gel probing of thick gels, in our characterization of GAPDH and actinin we used an electrophoretic probe delivery system to drive antibody from an agarose delivery gel into the polyacrylamide separation gel (again mitigating partitioning and achieving shorter assay times). This electrophoretic delivery system facilitated the use of standard primary/secondary antibody pairs, as described in the Methods. To expedite full-gel volumetric fluorescence readout of protein immunoblots, we employed light sheet microscopy in **Figure 5**. Comparison measurements using scanning laser confocal microscopy are presented in **Figure S4** (Supplementary Information). Data processing allows us to visualize immunoblot readouts as maximum intensity 3D renderings (**Figure 5**a), 2D contour plots showing X-Z fluorescence peaks for each fluorescence colour channel (corresponding to a target/antibody pair) (**Figure 5**b), and revolved 1D Z-directional fluorescence intensity plots (**Figure 5**c).

**Figure 5.**
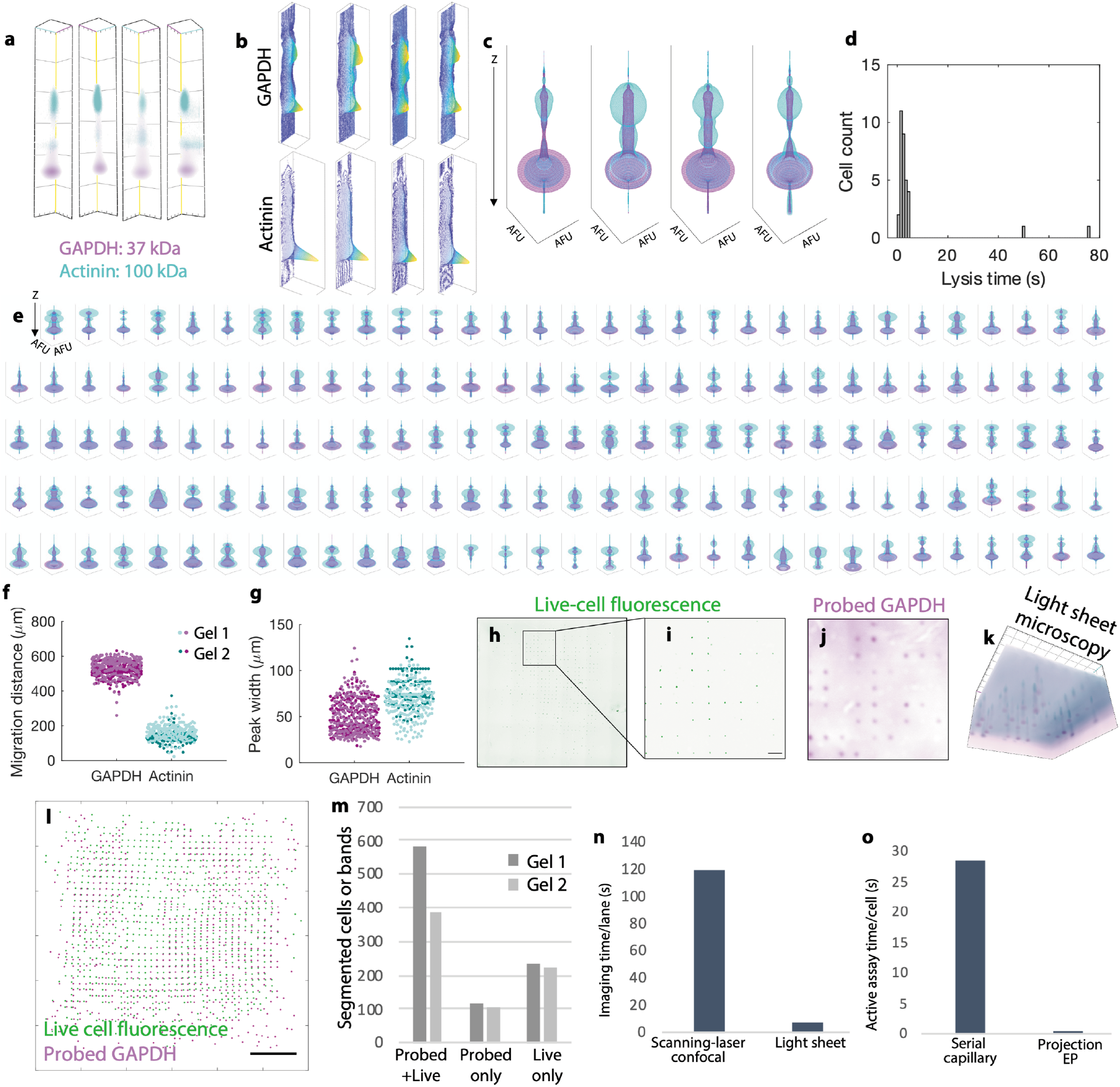
Projection electrophoresis permits simultaneous analysis of hundreds of single cells by concurrent separation after simultaneous lysis. **(a)** Maximum intensity projection 3D renderings of example separation lanes read out by tiled light sheet microscopy. **(b)** X-Z (Y-summed) contour plots of background-subtracted actinin and GAPDH protein signal within the lanes depicted in (a). **(c)** Revolved Z-intensity profiles for the four separation lanes depicted in (a-b). Each plot depicts Z-directional intensity profiles (summed fluorescence within each X-Y ROI vs. Z) for GAPDH (magenta) and actinin (cyan), revolved around the Z-axis to generate 3D rendering of fluorescence distribution. **(d)** Histogram quantification shows 86% of U251 cells lyse within 5 s of initiating lysis. **(e)** Quantified fluorescence intensity data for n=159 separation lanes passing quality control for both the GAPDH and actinin channels. Each plot depicts revolved Z-intensity profiles. **(f)** Quantified migration distances from a total of n=507 (GAPDH) and n=303 (actinin) lanes passing quality control in two projection electrophoresis gels. **(g)** Quantified Z-direction peak widths for the same bands analyzed in (f). **(h)** Full-gel wide-field fluorescence image of calcein-stained live BT474 breast tumour cells before analysis. **(i)** Subset of the live cells from (h), within a 1.75×1.75 mm light sheet microscopy field of view (scale bar depicts 200 μm). **(j)** Post-separation wide-field fluorescence image of probed GAPDH signal within the same field of view as (i). **(k)** Maximum intensity projection 3D rendering of a light sheet microscopy image (same field of view as (i-j)), showing 3D separations of GAPDH and actinin from tens of separation lanes, each corresponding to signal from the settled cells in microwells depicted in (h-i). **(l)** Overlay image of segmented spots corresponding to live BT474 cells in microwells (green) and probed GAPDH bands after separation (magenta), for the same separation gel depicted in (h-k). **(m)** Quantification of correspondence between the segmented live cells and bands (via intensity thresholding) within the same separation lanes as (l). **(n)** Per-lane imaging time of light sheet microscopy is >10X smaller than that for scanning laser confocal. **(o)** Projection electrophoresis demonstrates an 80X reduction in assay time per cell compared to serial capillary analysis.

Cells lyse nearly simultaneously (**Figure 5**d). Across 4 replicate lysis monitoring experiments, 31/36 monitored cells (86%) lysed within 5 s of placing the lysis gel on top of the microwell gel. Of the 5 remaining cells (all within the same replicate), 2 lysed during the 80 s monitoring period while 3 did not, potentially due to a bubble between the lysis and microwell separation gels. This near-simultaneous lysis, combined with parallelized electrophoretic separation over the full gel (>1000 separation lanes), facilitates concurrent analysis of hundreds of single cells. **Figure 5**e depicts revolved 1D intensity profiles for the 159 separation lanes within a single projection electrophoresis gel that passed R^2^ (>0.7 for Gaussian fit to 1D Z-intensity profile) and SNR (>3) quality control in both the actinin and GAPDH channels. Two hundred twenty-two separation lanes passed these quality controls in the GAPDH channel alone, while 204 lanes passed in the actinin channel alone. The intensity profiles show a GAPDH peak at a depth of 552±54 μm and an actinin peak at a depth of 165±42 μm (median ± one standard deviation). The profiles also show another peak in the actinin channel near the depth of the GAPDH peak, potentially due to off-target antibody binding and/or spectral bleed-through between the channels (optical filter sets) of the light sheet microscope. We observe the expected differential in electrophoretic mobility and nearly equivalent peak widths for the 37 kDa and 100 kDa targets for a total of n=507 (GAPDH) and n=303 (actinin) across duplicate separation gels (**Figure 5**f-g). Although from diffusion theory we would expect a larger peak width for the smaller protein target, differences in peak dispersion between targets can result in wider measured peak widths.

Projection electrophoresis is compatible with multi-modal imaging of the intact cells before separation, as well as the separated protein bands after the assay. Pre-separation live-cell imaging of intact BT474 cells (**Figure 5**h-i) correlated well with detected probed bands. The *in situ* separations facilitated by projection electrophoresis allow comparison of live-cell fluorescence – prior to projection immunoblotting (via wide-field fluorescence microscopy in **Figure 5**h-i) – and endpoint probed GAPDH signal (via wide-field fluorescence microscopy in **Figure 5**j and light sheet microscopy in **Figure 5**k). The analysis revealed appreciable spatial correlation between live cell imaging prior to separation and wide-field fluorescence images of separated GAPDH (**Figure 5**l-m). Comparison shows 63-74% of live cells detected are correlated with GAPDH detection. In two duplicate separations, 76% and 63% of detected live cells corresponded to probed GAPDH bands, 24% and 37% of detected live cells did not correspond to a probed GAPDH band, and 17% and 22% of probed bands did not visibly correspond to a live cell. This correlation (and potentially cell settling efficiencies and analysis throughput) could potentially be further improved in future work by encapsulating settled cells in hydrogel to mitigate cell loss/movement during manual gel transfer and electrophoresis stack setup.

We compared scanning laser confocal microscopy (**Figure S4** of the Supplementary Information) and light sheet microscopy (**Figure 5**) imaging of the same gel devices to acquire volumetric protein immunoblot readouts from the protein PAGE separation lanes. Each imaging modality presents a trade-off in field of view and Z-axis resolution, but the imaging throughput of light sheet microscopy was >10X higher than scanning laser confocal (**Figure 5**n), moving from ~120s/lane readout time down to ~8s/lane, while retaining sufficient Z-axis resolution to localize protein peaks. The laser scanning confocal imaging with 20X NA=1.0 water immersion objective (required for high-resolution optical sectioning) supported a 425 x 425 μm field of view, while light sheet microscopy with 5X detection objective (NA=0.16) provided a much larger 1.75 x 1.75 mm field of view. Because its optical sectioning is facilitated by the light sheet forming objectives forming a thin illumination sheet, light sheet microscopy allows the imaging optical section thickness to be decoupled from the detection objective NA, facilitating the use of lower NA detection objectives while maintaining optical sectioning^71^. The light sheet images acquired for projection electrophoresis had optical section thicknesses on the order of 10 μm, which should be sufficient to assess the diffusion-limited Z-directional peak widths of tens of microns for our separated protein bands.

Further, light sheet microscopy detected both protein targets with similar expected differential electrophoretic velocity and comparable peak widths of the immunoprobed targets to those measured with confocal (**Figure S4** of the Supplementary Information). With both readout modalities, we also observe the log-linear relationship between migration distance and molecular weight for endogenous targets that would be expected for a size-sieving separation (**Figure S5**). One consideration when comparing light sheet and confocal microscopy is refractive index mismatch between the gel and the objective immersion medium. In confocal microscopy, mismatch in sample vs. immersion medium refractive index results in distortion of the imaging point spread function, as well as distortion of the apparent scanned distance in z:

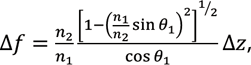

Where Δ*f* is the distance in z scanned by the focal spot, Δ*z* is the distance in z scanned by the objective, *n*_1_ is the objective immersion medium, *n*_2_ is the sample medium, and *θ*_1_ is the angle at which the marginal rays from the objective approach the top of the sample^72^. Although this equation derived from geometric optics is helpful in qualitatively understanding the effects of refractive index mismatch, experimental data often do not follow this relationship (likely due to the high sensitivity to *θ*_1_)^72^. In contrast, with light sheet microscopy we may not expect this distortion of apparent Z-directional distances because the position of the sectioned focal plane is controlled by the physical position of the light sheet within the sample, rather than by the optics of the detection objective. From this distortion function, we would expect to measure migration distances from confocal images shorter than those measured with light sheet microscopy (as n_2_, or the gel refractive index, is slightly higher than the water immersion medium n_1_). Comparing **Figure S4** of the Supplementary Information and **Figure 5**f, we indeed observe that confocal migration distances (**Figure S4**) are shorter than those measured from light sheet (**Figure 5**f). Given similar results in detection, migration location, and peak width for the model endogenous protein targets, the substantially larger field of view of light sheet microscopy proved beneficial, allowing endpoint imaging and analysis of 10x larger number of immunoblots (imaged separation lanes passing quality control: n=22 with confocal; n=303 actinin and n=507 GAPDH with light sheet microscopy, over two separation gels).

The parallel cell analysis approach described here overcomes shortcomings of serial analysis of cells. Although synchronous cell lysis is also not instantaneous across cells (with biological variation in lysis time on the order of seconds^73^), serial interrogation of individual cells leads to asynchronous analysis with longer time delays between analysis of the first cell and last cell in a population. Considering one example singlecell enzyme analysis separation, where individual cells are interrogated by a capillary sampler after cell lysis via a UV light pulse^17^, we estimate a 104 minute delay between interrogation of the first cell and interrogation of the last cell (219 cells analyzed with an analysis throughput of 2.1 cells/min). In contrast, concurrent analysis of 300 measurable separation lanes is completed with < 10 s delay between the first and last cell, assuming a small delay in cell lysis arising during the application of the lysis and solubilization buffer. By introducing a new, rapid, and parallelized electrophoresis approach, we demonstrate simultaneous single-cell separations of hundreds of single cells with an active assay time of 2.5 cells/s (from lysis through photocapture), depending on settling efficiency – this represents a >70-fold improvement in assay throughput over serial capillary systems as depicted in **Figure 5**o.

### Outlook

Projection electrophoresis addresses key bottlenecks in single-cell protein analysis by achieving rapid (<1.5 min active assay time), synchronous size-based protein separation for hundreds of unfixed cells in parallel, without sample transfer steps that can result in sample losses and changes in sample composition. Detection specificity combines antibody recognition with size-separation to confer proteoform-level specificity even when specific probes do not exist, and avoids the need for pre-separation tagging of proteins for detection. *In situ* cell lysis, separation, and photocapture of protein to the gel matrix mitigates deleterious effects of sample transfer between systems, including losses from adsorption to glassware, potential sample contamination, and sample changes between lysis and separation. Effectively synchronous analysis of hundreds of cells in parallel mitigates artifactual changes in cell population heterogeneity induced by heterogeneous lysis times, while also facilitating high assay throughput. Using rapid whole-cell lysis of unfixed cells, we demonstrate detection of both nuclear and cytosolic proteins. We design and characterize the system by both modelling and measuring the results of microscale physics. Compared with planar single-cell western blotting, we demonstrate a 10-fold reduction in sample volume to assay the same number of cells, and further throughput improvements may be possible by optimizing parameters for settling efficiency. Looking forward, the performance of projection electrophoresis can be improved in future work by moving to larger-area gels to further parallelize analysis, and by using our understanding of the driving small-scale physics to optimize gel materials for targets of interest and thus enable the use of even higher microwell densities and improved separation performance. With a straightforward, open microfluidic format and advantages complementary to existing protein analysis tools, we anticipate that projection electrophoresis will assist in the development of proteoform-level atlases of single-cell diversity.

## Materials and Methods

### Polyacrylamide gel fabrication

Substrate-free and featureless polyacrylamide gels (used for lysis and enclosing purified protein microwells) were fabricated between a glass plate (McMaster-Carr) and 25×75 mm or 50×75 mm glass slide (VWR), both treated with Gel Slick^®^ glass plate coating according to the manufacturer’s instructions and rinsed briefly in deionized water prior to gel fabrication. For purified protein experiments and cell seeding gels, microwell-stippled gels were cast between a methacrylate-functionalized glass slide (treated to promote gel adhesion as previously described by our group^59^) and a dichlorodimethylsilane (Sigma Aldrich 440272)-treated silicon wafer (WaferPro). The wafer was patterned with SU-8 3050 (Microchem) features and was silane-treated to facilitate gel release as previously described by our group^74^. Wafers with 40 μm high SU-8 features (32 μm diameter microwells for purified protein experiments and lysis monitoring, or 25 μm diameter microwells for BT474 and U251 single-cell separations) were fabricated according to the manufacturer’s instructions and then silane-treated as previously described^59^. Our approach for substrate-free micropatterned gel fabrication was inspired by that recently developed by our group to fabricate releasable gel microparticles^74^. In all cases, 1 mm or 500 μm thick gels were fabricated using spacers of the appropriate thickness (C.B.S Scientific Gel Wrap) between the two glass pieces or between the wafer and glass slide.

Acrylamide precursor solutions for the various gels were prepared by diluting 30% stock acrylamide/bis-acrylamide precursor (Sigma-Aldrich A3699) and Rhinohide^®^ (ThermoFisher R33400) solution (to increase mechanical robustness of substrate-free gels) in ultrapure water (Millipore^®^) and 10X tris-glycine (Bio-Rad 1610734) where appropriate. Separation gels contained 3 mM final concentration BPMA: N-(3-((3-benzoylphenyl)formamido)propyl) methacrylamide, which was custom-synthesized by PharmAgra Labs (cat. no. PAL0603)^59^. BPMA is copolymerized into the gel matrix and permits photo-immobilization of proteins. Gels were chemically polymerized for 60 minutes with 0.08% (w/v) ammonium persulfate (APS, Sigma-Aldrich A3678) and 0.08% (v/v) TEMED (Sigma-Aldrich T9281), from freshly-prepared 10% stock solutions in ultrapure water. The constituents of the various gel types used in this work are presented in Supplementary Table S1, and a schematic showing the released gel fabrication and molding process is depicted in **Figure S6** (Supplementary Information).

After polymerization, gels were trimmed to size using a razor blade and released from the glass substrate by carefully sliding the razor blade under the gel, applying firm pressure with the blade onto the glass and gently adding water between the razor blade and gel to lubricate and prevent tearing. Separation and shield/lysis gels were stored in the appropriate buffer solution for a minimum of 12 hours and up to 4 days prior to running separations. Buffer storage conditions are presented in Supplementary Table S1 and were either (1) dual function lysis-electrophoresis modified RIPA buffer (1X or 2X RIPA as described in Supplementary Table S2), (2) run buffer (1X tris-glycine with 0.5% Triton X-100), or (3) 1X tris-glycine for gels to be used for cell settling. Buffers used in this work are described in Supplementary Table S2.

### Z-directional electrode system

The Z-directional electrode separation system consists of planar electrodes integrated into a custom laser-fabricated acrylic alignment setup and brought into contact with two 32 mm diameter, 3 mm thick neodymium rare-earth magnets on the back side of each electrode (each magnet specified to provide 19 lbs of pull force). Uniform spacing between the electrodes is facilitated by 2.5 mm (purified protein experiments) or 3 mm (cell experiments) removable polymer spacers (C.B.S Scientific Gel Wrap) at the top and bottom of the electrodes. The planar electrodes were commercial platinum-coated electrotransfer anodes (Bio-Rad Criterion anode plates) with plastic housings modified to permit close proximity of the electrode surfaces. Electric fields were provided by a power supply (Bio-Rad PowerPac^®^ Basic) connected to the electrodes with standard banana plug interfacing. Cold packs on the back side of each electrode maintained gel temperature at ~4°C to help to mitigate deleterious effects of Joule heating during purified protein separations. To aid in lysis and protein solubilization^60^, the cold packs were heated in a 55°C water bath for single-cell separations, yielding electrode temperatures of ~37°C. Hot packs were heated for >10 minutes to equilibrate to temperature, and exchanged every ~15 minutes between separations.

### Purified protein separation experiments

Mixed molecular weight purified protein solutions were prepared by diluting stock solutions of Alexa Fluor^®^555-labelled bovine serum albumin (BSA; Thermo Fisher Scientific A34786; 5 mg/mL stock), Alexa Fluor^®^488-labelled ovalbumin (OVA; Thermo Fisher Scientific O34781; 2 mg/mL stock), and Alexa Fluor^®^647-labelled donkey anti-mouse secondary antibody (IgG; Thermo Fisher Scientific A31571 lot 1900251; 2 mg/mL stock) in a run buffer (Supplementary Table S2) consisting of 1X tris-glycine (prepared by ultrapure water dilution of 10X stock, Bio-Rad 1610734) containing 0.5% v/v Triton X-100 (Sigma). High molecular weight purified protein solutions were prepared by diluting stock solutions of Alexa Fluor^®^488-labelled lectin (Thermo Fisher Scientific L11270; 2 mg/mL stock), and Alexa Fluor^®^647-labelled transferrin (Thermo Fisher Scientific T23366; 5 mg/mL stock) in the same run buffer. All proteins were diluted to a final concentration of 5 μM.

Western blotting filter paper of 1 mm thickness (Thermo Fisher Scientific 84783) was cut into 12×12 mm squares and allowed to equilibrate in dual-function lysis-electrophoresis modified RIPA buffer for >10 minutes prior to starting separations. Microwell-patterned separation gels of 1 mm thickness and shield gels of 500 μm thickness were prepared as described in Supplementary Table S1, cut into squares of 9×9 mm (separation gel) or ~12×12 mm (shield gel), and equilibrated in the appropriate buffer for >12h.

We first assessed the appropriate constant current conditions to yield the target electric field. We set up a dummy separation stack consisting of the anode (bottom), the buffer-equilibrated filter paper, a separation gel, a shield gel (with bottom surface dried by placing on a clean, dry Kimwipe), and the cathode (top). We supplied a constant voltage of 13 V (the necessary voltage for an electric field of 52 V/cm) and noted the initial current through the dummy stack (typically ~33 mA). This constant current was chosen for each following separation run on a given day, and the initial and final voltages were noted during each trial to quantify the electric field and resistance changes during the separation.

To run the separations, we again stacked the anode, buffer-soaked filter paper, and separation gel, but this time dried the top (microwell-studded) surface of the separation gel gently bringing it into contact with a folded Kimwipe prior to stacking on top of the filter paper. Drying the top surface of the separation gel and bottom surface of the shield gel reduces dilution of the purified protein solution prior to separation. We pipetted 3 μL of the mixed purified protein solution (either the mixed molecular weight standard or the high molecular weight standard) on top of the separation gel and spread the resulting droplet to cover the surface of the gel using the side of a p20 pipette tip, taking care not to puncture the gel surface. We then dried the bottom side of a shield gel by placing it on a folded Kimwipe, brought it into contact with the separation gel by carefully lowering it from one corner to reduce bubble entrapment, and assembled the cathode on top. We supplied constant current between the anode and cathode for varying electrophoresis times, running duplicate gels for each electrophoresis time in each experiment.

Immediately after electrophoresis was complete, the power was shut off and the system disassembled to permit optical access for a UV source (Hamamatsu Lightningcure LC5). The liquid light guide-coupled UV source was used to photocapture the separated protein bands using a 45s UV exposure, holding the tip of the liquid light guide ~4 cm from the separation gel. We have photocaptured with both the top (microwell) and bottom (flat) side of the separation gel facing the light guide. The total time between beginning electrophoresis and initiating UV exposure was recorded for each test as an estimate of the in-gel diffusion time *t_diff_*.

After photocapture, each gel was rinsed briefly in deionized water and then stored for >12 hours in 1X tris-buffered saline solution with Tween^®^ (TBST, prepared from Cell Signaling Technology 9997S 10X stock and MilliQ ultrapure water) in a polystyrene 12-well plate prior to imaging to permit release of any non-photocaptured protein. Gels were imaged through a #1 coverslip using a Zeiss LSM 880 laser-scanning confocal microscope fitted with a 20X water dipping objective (NA=1.0, Zeiss W Plan APO 20x/1 DICIII). A confocal Z-slice spacing of 5 μm was chosen, and volumes extending ~100 μm past visible fluorophore bands were imaged. As we were not quantifying or comparing protein abundance, excitation laser powers were adjusted to permit fluorescence visibility depending on the sample brightness. Fluorescence intensities were not compared between purified protein samples. Similarly, images were brightness and contrast-adjusted in Fiji^75^ (based on ImageJ^76^, National Institutes of Health) to ensure visibility of protein bands. Maximum intensity projection 3D renderings were prepared in Olympus Zen Blue software.

Purified protein confocal datasets were analyzed using custom analysis scripts in MATLAB^®^. The scripts were designed to (1) find and track regions of interest (ROIs) corresponding to protein originating from each of the microwells through the depth of each confocal stack, (2) create 1D intensity plots of summed fluorescence intensity vs. Z depth for each ROI by summing the fluorescence intensity for the ROI at each Z-plane, (3) assess the Z migration distance and peak width for each protein by Gaussian fitting each intensity profile peak, allowing the user to set bounds for fitting peaks corresponding to each purified protein, (4) measure the diffusional spreading of protein from each microwell by Gaussian fitting the summed 1D X- and Y-intensity profiles for each protein at its Z migration peak location for each protein, (5) plot migration distance, Z-direction peak width, and X-Y peak width vs. electrophoresis time or diffusion time, comparing across multiple gels.

Zeiss CZI confocal Z-stacks and associated metadata were imported into MATLAB^®^ (MathWorks^®^) using the MATLAB^®^ Bio-Formats libraries provided by the Open Microscopy Environment^77^. ROIs were segmented in each Z-slice image (summing the intensities of all colour channels into one image for the purposes of segmentation) using intensity thresholding followed by morphologic open and close operations to remove erroneously-segmented small features and close incomplete contours. A fill operation was then used to close all holes in the segmented spots of protein, and all segmented objects touching the border of the image were removed. The centroids of the segmented spots of protein at each Z location were stored, and a MATLAB^®^ particle tracking library (based on a previous IDL implementation^78^) made publicly accessible by Prof. Daniel Blair and Prof. Eric Defresne^79^ was used to track the positions of protein originating from each microwell through the Z depth of the image. The tracked centroids were then subject to a quality control step to remove protein spots that were only found in small portions of the full Z volume. After this was complete, the tracking code output a set of tracked ‘particles’ (protein spots originating from a given microwell), each with a list containing the x-y location of the centroid of each protein spot for every Z location in the image. To create the intensity profiles, the intensities in a 300 pixel (102 μm) square ROI were analyzed surrounding each centroid at each Z location. The data were background-subtracted by subtracting from each pixel the average measured intensity 15 μm below the bottom of the microwells, in regions at least 100 pixels (34 μm) from any segmented protein spots. After Gaussian fits to find the migration distance, z-direction peak width, and x-y peak width for each protein peak, the data from all of the ROIs from multiple gels (multiple electrophoresis times in duplicate) were plotted and fit to the expected linear physical relationship (migration distance vs. electrophoresis time, squared peak width and squared x-y peak width vs. total in-gel time).

### Cell culture

U251 human glioblastoma cells stably transduced with turboGFP by lentiviral infection were kindly provided by Prof. Sanjay Kumar’s laboratory at UC Berkeley. Cells were maintained in tissue culture flasks in a standard cell culture incubator (Heracell 150i) at 5% CO2 and 37°C, in DMEM (Invitrogen 10566016) supplemented with 10% FBS (Gemini Bio-Products Benchmark 100-106), 1% penicillin/streptomycin (Life Technologies 15140-122), 1X sodium pyruvate (Thermo Fisher 11360070), and 1X non-essential amino acids (Life Technologies 11140-050 100X stock). The cells were passaged at a density of 1:10 to 1:40 after reaching ~80% confluency by detaching with 0.05% Trypsin-EDTA (Life Technologies 25300120), centrifuging for 3 minutes at 1000 RPM, and resuspending in completed media to reseed.

BT474 human breast cancer cells were purchased from the UC Berkeley Biosciences Divisional Services Cell Culture Facility. Cells were maintained in tissue culture flasks in a standard cell culture incubator (Heracell 150i) at 5% CO2 and 37°C, in DMEM (Invitrogen 10566016) supplemented with 10% FBS (Gemini Bio-Products Benchmark 100-106) and 1% penicillin/streptomycin (Life Technologies 15140-122). The cells were passaged at a density of 1:2 to 1:8 after reaching ~80% confluency by detaching with 5 mM EDTA in PBS (Invitrogen 15575-038 diluted in sterile 1X PBS: Life Technologies 10010049), centrifuging for 3 minutes at 1000 RPM, and resuspending in completed media to reseed.

### Cell lysis monitoring experiments

We compared diffusion profiles of cells lysed after settling in 32 μm diameter, 40 μm high microwells within 1 mm thick gels. Microwell gels and lysis gels (18×18 mm in area) were prepared and equilibrated in PBS for >12h as described above.

To settle cells in microwells, we followed a procedure similar to that used for single-cell western blotting^59^: U251-turboGFP cells were detached using 0.05% trypsin-EDTA (Life Technologies 25300120), resuspended in PBS (Life Technologies 10010049) at a concentration of 100 000 cells/mL, and filtered using a cell strainer (Corning 352235). The 18×18 mm microwell gels were placed microwell side up in a 60 mm tissue culture dish. ~100 μL cell solution was pipetted onto the surface of each microwell gel (to cover but avoid spillage over the edge of the gel) and spread to cover the full array of microwells. The cells were allowed to settle for 5 minutes on ice, gently agitating every ~2 minutes, before rinsing with 1-3 mL PBS by tilting the tissue culture dish at a ~40° angle, pipetting PBS at the top surface of the microwell gel, and allowing it to flow over the surface of the microwell gel into the bottom of the petri dish. After rinsing was complete, the excess PBS/cell solution was aspirated for biohazard disposal.

To facilitate monitoring, each microwell gel was immobilized within a 60 mm petri dish by pipetting 200 μL of a warmed solution of 5% agarose (Invitrogen 16500) in PBS (Life Technologies 10010049) beside the gel and allowing it to gel at room temperature (in contact with the edge of the gel and the petri dish). Before lysis, excess fluid was removed from the gel by tilting the petri dish and wicking away the fluid layer by bringing a folded Kimwipe into contact with the corner of the gel (not touching cell-containing regions). The petri dish was then secured to the stage of an Olympus IX71 microscope for monitoring with a 4X or 10X air objective. Fluorescence excitation was provided by an X-Cite source (Excelitas Technologies) through a GFP filter set (Chroma 49011 ET), and fluorescence was measured using an EM-CCD camera (Andor iXon). Time-lapse images of the turboGFP fluorescence were captured using the MetaMorph^®^ imaging software (Molecular Devices). After focusing, setting up the imaging settings and initiating the time-lapse, the lysis gel was carefully placed on top of the cell-containing gel, starting with one corner and then smoothly bringing the rest in contact to reduce bubble entrapment between the two gels. The time at which the lysis gel was placed in the time-lapse series was recorded. The lysis gel was a 20%T, 10% Rhinohide shield gel equilibrated in 2X RIPA-like lysis buffer.

The lysis monitoring data were analyzed using custom scripts written in MATLAB^®^. At each time point, the cells were segmented using adaptive thresholding of the median filtered (3×3 neighborhood) image. The segmentation was improved using morphologic open and close operations, and the centroids of the segmented cells were computed and stored. The MATLAB^®^ particle tracking library (based on a previous IDL implementation^78^) made publicly accessible by Prof. Daniel Blair and Prof. Eric Defresne^79^ was used to track the centroids of each cell over the course of the experiment, although minimal drift was observed. The maximum fluorescence intensity and total fluorescence intensity in a 100 μm diameter circle surrounding the centroid of each segmented cell were tracked for each time point. All pixel intensities were background-subtracted using the average fluorescent intensity of the background region at each time point, taken to be the image regions greater than 30 μm away from all segmented regions.

### Finite-element modelling of protein diffusion

We used finite-element modelling of dilute species transport in COMSOL^®^ Multiphysics to predict protein concentrations during lysis and electrophoresis. The simulation geometries for the Z-directional simulations are presented in the cross-sectional view shown in **Figure 4**b. 2D axisymmetric models were used for all simulations due to the inherent symmetry of the geometry. Diffusion coefficients in free solution portions of the model were estimated from the Stokes-Einstein equation^49^, while in-gel diffusion was estimated from the free-solution diffusivity using the methods presented by Park, *et al*.^50^ Protein hydrodynamic radii were estimated from the number of amino acids^80^. Thermodynamic partitioning of protein was simulated using flux boundary conditions relating in-gel concentration to in-solution concentration using a partition coefficient *k=C_gel_/C_solution_*. Partition coefficients were estimated using the Ogsten model^81^ (assuming size-exclusion partitioning), using estimates of fibre radius from Tong and Anderson^65^ and hydrogel volume fraction from the data presented by Baselga et al^82^. A temperature of 4°C was assumed for all simulations except the comparison with the single-cell western blot platform, which assumed 10°C for both platforms. The diffusion and partition coefficients for each protein at 4°C are presented in Table 1.

In each model, gel-solution boundaries were modelled as flux boundary conditions taking size-exclusion partitioning into account. The edges of each model were modelled as flux boundary conditions permitting protein to freely leave the model; however, the simulation region was sufficiently large (500×500 μm) that changing these boundary conditions to no flux resulted in negligible change to the modelled concentration profiles. The initial protein concentration in the cell region was 2 μM, while the initial concentration elsewhere in the model was zero. In models of the single-cell western blotting platform, the bottom surface of the gel was modelled as no flux to model the presence of the glass slide present in that system. The model was meshed with a physics-controlled mesh calibrated for fluid dynamics, and a user-controlled override with maximum element size of 0.5 μm was used in the microwell and thin fluid layer regions to ensure sufficient mesh density in these smaller regions.

A time-dependent study was used to model the protein concentration profile during lysis and electrophoresis. To model both in the same diffusive model, the diffusion and partition coefficients were set as step functions. For the first 25s of the model (the lysis portion), the diffusion and partition coefficients were set as in Table 1. After this 25s lysis period, all partition coefficients were set to 1, and the diffusion coefficients in all regions of the model were set to those for 7%T gel (the separation gel), effectively simulating instantaneous injection of the full protein profile into a separation gel. While this method is straightforward and does not require modelling of the electrophoresis physics, it provides a conservative (over-) estimate of Z-directional diffusional spreading because it does not model stacking of the protein band as it is injected from free solution into the gel.

After running the model, we assessed the maximum protein concentrations in the simulated geometry. We also assessed the integrated protein intensities (the protein concentrations at each radial (x) location, integrated in z) to model the wide-field microscopy imaging measurements in which fluorescence from the full Z region is integrated into a 2D image. These intensities were compared with the experimental lysis monitoring data.

### Single-cell separations

Adherent U251 glioblastoma and BT474 breast tumor cells were detached from culture flasks with 0.05% Trypsin-EDTA (Life Technologies 25300120) (U251) or 5 mM EDTA (Invitrogen 15575-038) in PBS (BT474) and resuspended in cold PBS (Life Technologies 10010049) at a concentration of 1.5×10^6^ cells/mL. For viability staining to aid in microwell occupancy assessment, BT474 cells were stained with Calcein-AM in incomplete DMEM for 20 minutes at room temperature prior to resuspension. Cell suspensions were kept on ice and filtered through a cell strainer (Corning 352235) to reduce cell aggregates immediately prior to settling.

1 mm-thick micropatterned separation gels were stored in 1X tris-glycine (Bio-Rad) and buffer-exchanged to lower conductivity sucrose-dextrose dielectrophoresis buffer (DEP buffer: 2.39 g/L HEPES (VWR 3638C017), 80.7 g/L sucrose (Sigma-Aldrich S0389), 4.5 g/L dextrose (Fisher D16-500), 11.1 mg/L CaCl2 (FisherC79-500); pH adjusted to pH 7.5 with NaOH)^83^ at least 10 minutes prior to cell settling. To settle the cells, gels were placed microwell side up inside of a 35 mm tissue culture dish. 25 μL of single-cell suspension was supplied to each 9×9 mm gel (by first pipetting 15 μL onto the gel surface, spreading with a pipette tip while taking care not to perforate the gel surface, and subsequently dispensing another 10 μL onto the surface. Cells were allowed to gravitationally settle for 20 minutes, agitating the gel periodically and covering the gel with the lid of the 35 mm tissue culture dish (to reduce evaporation). Settled cells were checked at the 10-minute mark, and if the cell suspension had aggregated towards the centre of the gel, an additional 10 μL of cell suspension was supplied to the gel edges.

After 20 minutes of settling, gels were rinsed by holding the petri dish at a ~40° angle and pipetting 0.5 mL of DEP buffer onto the top corner of the gel, allowing the fluid stream to wash over the full gel into the tissue culture dish. The wash fluid was aspirated for biohazardous waste disposal, and the wash was repeated with an additional 0.5 mL of DEP buffer before pipetting ~40 μL DEP buffer on top of the gel to prevent drying during imaging. Tiled images of the settled cell fluorescence were captured using the ScanSlide plugin for the MetaMorph^®^ imaging software (Molecular Devices), using an Olympus IX51 inverted wide-field fluorescence microscope fitted with an X-Cite^®^ illumination source (Excelitas Technologies), GFP filter set (Chroma 49011 ET), 4X objective, and CoolSNAP HQ2 CCD camera (Teledyne Photometrics). Cell occupancy was calculated from a MATLAB^®^ script that analyzed tiled live-cell brightfield and fluorescence images, determined whether there was a fluorescent object in a region surrounding each microwell (via thresholding segmentation) and, if found, displayed the region and prompted user input of number of cells in the microwell region.

Aliquots of lysis buffer (4 mL) were prepared by dissolving urea to a final concentration of 8 M in 2X RIPA-like lysis buffer in a water bath set to 55°C. Lysis gels (14×14×1 mm) were stored in 2X RIPA-like lysis buffer and transferred to aliquots of urea lysis buffer as soon as the urea had dissolved (10-60 minutes prior to running the separations). The lysis gels in buffer aliquots were heated to 55°C in a water bath until immediately prior to use, agitating periodically to ensure the solution was well mixed.

After cell settling and live-cell imaging, 4 mL 2X tris-glycine (diluted from Bio-Rad 1610734 10X stock) was pipetted into the tissue culture dish containing the separation gel and incubated for 1 minute to reduce the concentrations of potentially unwanted ions. A 10×10×1 mm filter paper (cut to size from Thermo Fisher Scientific western blotting filter paper) was hydrated in the heated lysis buffer aliquot and placed on the anode. The separation gel was placed (microwell side up) on top of the filter paper immediately after tris-glycine incubation, placing slowly from one corner to reduce bubble entrapment between the layers. The lysis gel was removed from the buffer aliquot and placed on top of the separation gel, again placing gradually so as to not introduce bubbles between the gels. The lysis timer was started as soon as the lysis gel was placed, and the electrode system was closed by placing the cathode on top of the lysis gel. After 25s lysis, the separation was initiated by supplying 80 mA of constant current (typically yielding 13-16V initial voltage for an average electric field of 43-53 V/cm across the gel stack) using a DC power supply (Bio-Rad PowerPac Basic) and recording the voltage at 5s intervals during electrophoresis. After electrophoresis was complete, the power supply was stopped, electrode system opened, and 45s photocapture was completed using a Hamamatsu Lightningcure LC5 UV source. The gel was then rinsed briefly in deionized water before equilibrating in tris-buffered saline with Tween^®^ (1X TBST, from Cell Signaling Technology 9997S 10X stock) overnight to remove any residual lysis buffer (exchanging the buffer after 2h). Projection electrophoresis gels probed with F(ab) fragments were blocked overnight in 5% BSA in TBST at 4°C or for between 2-4h on a shaker at room temperature prior to immunoprobing.

### Immunoprobing and imaging

Single-cell projection electrophoresis immunoblotting gels were probed either diffusively (for initial characterization experiments using labelled F(ab) fragments in **Figure 4**) or electrophoretically (using standard primary and fluorescently-labelled secondary antibodies in **Figure 1** and **Figure 5**). All probing wash steps were performed using an electrophoretic wash platform.

In-gel probing requires high concentrations of immunoprobes to mitigate size-exclusion partitioning effects^59,65^. To minimize reagent consumption, we probe single-cell western blotting gels using minimal solution volumes. For the projection electrophoresis system, larger probe volumes are required to probe thicker gels; as such, we deliver probes using thin (0.5-1 mm thick) hydrogel layers to provide even probe delivery to all regions of the gel. Fluid, in contrast, tends to pool around gel edges, resulting in brighter probed signal at the gel edge. Constituents and fabrication parameters of probe delivery gels are described in Table 2.

**Table 2.** Probe delivery gel fabrication parameters for electrophoretic and diffusive probing.

| Gel type | Final agarose concentration after probe dilution | Antibody probes | Gelation time | Fabrication setup |
| --- | --- | --- | --- | --- |
| Primary antibody agarose electrophoretic probe delivery gel | 1.5% g/ml (dissolved in 1X tris-glycine) UltraPure LMP Agarose, (Thermo-Fisher: 16520050) | 10% v/v final from stock of each: (1) Rb α-actinin & (2) Gt α-GAPDH or (1) Ms α-PTBP1 & (2) Gt α-GAPDH | 5 mins on ice pack | Glass plate and glass slide |
| Secondary antibody agarose electrophoretic probe delivery gel | 1.5% g/ml (dissolved in 1X tris-glycine) | 10% v/v final from stock of each: (1) Dk α-Rb AF555 & (2) Dk α-Gt AF488 or (1) Dk α-Ms AF555 & (2) Dk α-Gt AF488 | 5 mins on ice pack | Glass plate and glass slide |
| hFAB agarose diffusive probe delivery gel | 1.5% g/ml - dissolved in 1X tris-buffered saline solution with Tween® (1X TBST, from Cell Signaling Technology 9997S 10X stock) | 10% v/v final from stock of either: (1) Bio-Rad hFAB Rhodamine α-GAPDH or (2) hFAB Rhodamine α-tubulin | 5 mins on ice pack | Glass plate and glass slide |

For diffusive probe delivery, Bio-Rad hFAB probes for GAPDH (12004167) or tubulin (12004165) were mixed at a 1:10 dilution with low melting temperature agarose (Invitrogen 16520-050) solution. Agarose was dissolved in 1X tris-buffered saline solution with Tween^®^ (TBST) to yield a final concentration of 1.5% (w/vol) after probe dilution, and maintained on a hotplate with spin bar at a temperature of ~40°C until mixing with the probes. Temperature of the solution immediately after mixing typically read ~30°C. The agarose gel was then immediately cast from the agarose-probe solution by pipetting between a heated (to ~30°C) glass plate and microscope glass slide, separated by gel casting spacers (C.B.S. Scientific GelWrap, 0.5 mm thickness). After casting, the glass plate setup was carefully moved onto a cold pack and the agarose was permitted to gel for 5 minutes before carefully disassembling the stack, cutting the gels to match the size of the separation gels, and immediately setting up the probing stacks. Each high-density separation gel was sandwiched between two 0.5 mm thick agarose probe delivery gels in a 24-well plate (with surrounding wells filled with water), sealed with a plate sealer, and incubated at 4°C in the dark for 40 hours.

For electrophoretic probe delivery, agarose probe delivery gels were fabricated in the same manner as the diffusive probe delivery gels described above; however, gels were fabricated at 1 mm thickness. Primary antibodies were electrophoretically introduced, incubated, and electrophoretically washed; this process was then repeated for fluorescently-labelled secondary antibodies. Primary antibodies used were Rb α-actinin (CST 6487, lot 2), Ms α-PTBP1 (Sigma WH0005725M1, lot J4241-3H8), and Gt α-GAPDH (SAB2500450, lot 6377C3); secondary antibodies were Dk α-Rb AF555 (A-31572, lot 2017396), Dk α-Ms AF555 (A-31570, lot 2045336), and Dk α-Gt AF488 (A-11055, lot 2059218). Immediately after casting the antibody probe delivery gel, a stack was set up for electrophoretic probe introduction. The probe delivery gel was placed against the flat (non-microwell-stippled) side of the separation gel. The stacked gel setup was sandwiched between two pieces of western blotting filter paper (Thermo Fisher Scientific 84783) held together using a laser-cut acrylic holder with a cut-out for the gel stacks. The holder was used to suspend the gels (with the separation gel facing the anode (+) and delivery gel facing the cathode (-)) in a slab-gel blotting module (Invitrogen X-Cell II) filled with 1X tris-glycine with 0.5% Triton X-100. Probes were transferred from the delivery gel into the separation gel at an electric field of 8 V/cm for 13 minutes; the stack was then disassembled and the separation gels were incubated at room temperature on a glass slide within a hydration chamber for 2 hours for primary antibody binding, or 1 hour for secondary antibody binding.

After each probe incubation step (primary and secondary antibodies), projection electrophoresis gels were electrophoretically washed by sandwiching between two filter papers soaked in 1X tris-glycine with 0.5% Triton X-100 (held in place by custom laser-cut acrylic with a cutout for the gels, which suspended the gels within the chamber of a slab-gel blotting module. Separation gels were submerged in 1X trisglycine with 0.5% Triton X-100 for ~1 min for rehydration immediately before electrophoretic washing. The blotting module was also filled with 1X tris-glycine with 0.5% Triton X-100. Gels were electrophoretically washed for 15 minutes at an electric field of 12 V/cm.

Gels were confocal imaged through a #1.5 coverslip using a Zeiss LSM 880 laser-scanning confocal microscope fitted with a 20X water dipping objective (NA=1.0, Zeiss W Plan APO 20x/1 DICIII). A confocal Z-slice spacing of 5 μm was chosen, and volumes extending ~100 μm past visible fluorophore bands were imaged. As we were not quantifying or comparing protein abundance, excitation laser powers were adjusted to permit fluorescence visibility depending on the sample brightness, as fluorescence intensities were not compared between cell separations. Similarly, images were brightness and contrast-adjusted in Fiji^75^ (based on ImageJ^76^, National Institutes of Health) to ensure visibility of protein bands.

For full-gel imaging, gels were imaged using a Zeiss Lightsheet Z.1 system fitted with a 5X detection objective (Zeiss EC Plan-Neofluar 420330-8210) and 4X light sheet forming objectives (Zeiss LSFM 400900-9010). Samples were excited with 488 nm and 561 nm lasers and detected using two pco.edge sCMOS cameras (with filter sets for AlexaFluor^®^ 488 and AlexaFluor^®^ 555). The samples were mounted to #1.5 coverslips using superglue at the gel corners, and the coverslip was glued to a custom 3D printed adapter to suspend the gel within the imaging chamber. The gels were imaged facing the detection objective (not imaged through the mounting coverslip). Tiled Z-stack images with 5-6 μm Z-slice spacing were acquired over the full gel volume, with 10% overlap between acquisition fields of view to ensure complete gel coverage. 49-56 Z-stack fields of view were typically required to cover the full gel area, each with 200-300 Z-slices.

Confocal and light sheet microscopy datasets were analyzed using custom analysis scripts in MATLAB^®^, similar to those described above for purified protein datasets. Zeiss CZI confocal Z-stacks and associated metadata were imported into MATLAB^®^ (MathWorks^®^) using the MATLAB^®^ Bio-Formats libraries provided by the Open Microscopy Environment^77^. The analysis workflow is described in Figure S7. 3D datasets are made up of 2D (X-Y) slice imaged that are each processed to assess the 3D positional data for each separated protein peak). Intensity profiles were background-subtracted by subtracting the average intensity of a 5-pixel (light sheet) or 20-pixel (confocal) border surrounding each X-Y ROI from each ROI pixel at each Z location prior to summing the intensities to generate Z-intensity profiles. Thin (10-pixel for confocal and 5-pixel for light sheet) borders between the ROI and background regions were used to assess the background-subtracted noise of the measurement to compute signal-to-noise ratios for each protein peak. Protein peaks corresponding to separation lanes were quantified (passing quality control) if: (1) there was a segmented fluorescence spot, (2) the Gaussian fit to the Z-intensity profile of the peak region had an R^2^ of the Gaussian intensity profile fit of >0.7, and (3) the Gaussian fit had a signal-to-noise ratio (Gaussian peak fit amplitude divided by twice the standard deviation of the background-subtracted noise calculation region over the Z region of the protein peak) of >3. A schematic representing the analysis workflow for tiled light sheet images (similar to the workflow used for confocal images but without the tiled functionality) is presented in Figure S8 of the Supplementary Information.

## Supporting information

Electronic Supplementary Information

## Acknowledgements

This work was funded in part by the National Cancer Institute of the National Institutes of Health, Cancer Moonshot award, Grant Number: 1R33CA225296-01 to A.E.H, and by the Chan Zuckerberg Biohub. S.M.G. gratefully acknowledges the support of a postdoctoral fellowship from the Natural Sciences and Engineering Research Council of Canada (NSERC). A.P.M. gratefully acknowledges support from the National Science Foundation Graduate Research Fellowship Program (NSF GRFP). We sincerely thank Dr. Burcu Gümüscü Sefünc for her assistance with early proof-of-concept tests. Confocal and light sheet imaging experiments were conducted at the CRL Molecular Imaging Center at UC Berkeley, supported by the Helen Wills Neuroscience Institute. We acknowledge all members of the Herr Lab at UC Berkeley, as well as Dr. Ben Smith at UC Berkeley and the group of Prof. Mark Pegram at Stanford University, for useful discussions and feedback. We are also grateful to the CRL Molecular Imaging Center staff as well as Kamran Ahmed and Eva Nichols for assistance with light sheet imaging setup.

## Author contributions

S.M.G. and A.E.H. designed the projection electrophoresis assay. S.M.G., A.P.M., and A.E.H. designed experiments. S.M.G. performed projection electrophoretic separations of purified proteins and single cells. S.M.G. and A.P.M. performed electrophoretic probing experiments. S.M.G. designed software and performed data analysis. All authors wrote the manuscript.

