## Supplementary material for "3D projection electrophoresis for single-cell immunoblotting": Electronic Supplementary Information

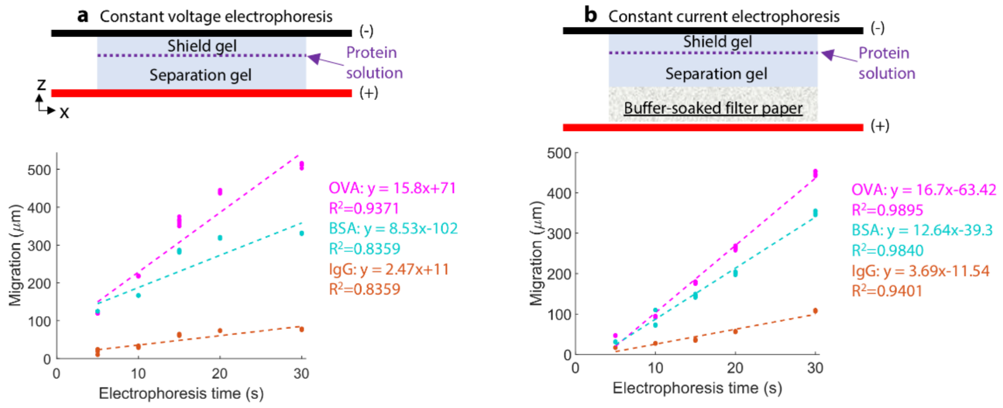


**Figure S1**. Optimization of Z-directional electrophoresis system to facilitate constant-velocity migration. **(a)** Shows a cross-sectional view of the setup before optimization. The separation gel is in direct contact with the anode, and the electric field was supplied as constant voltage. Linear fits to the migration data are poor, with migration slowing at increasing electrophoresis times. **(b)** Depicts the system after optimization. A buffer-soaked filter paper is placed between the separation gel and the anode to mitigate pH changes due to electrolysis at the electrode surface, and the electric field was supplied as constant current. Linear fits to the migration data are improved.

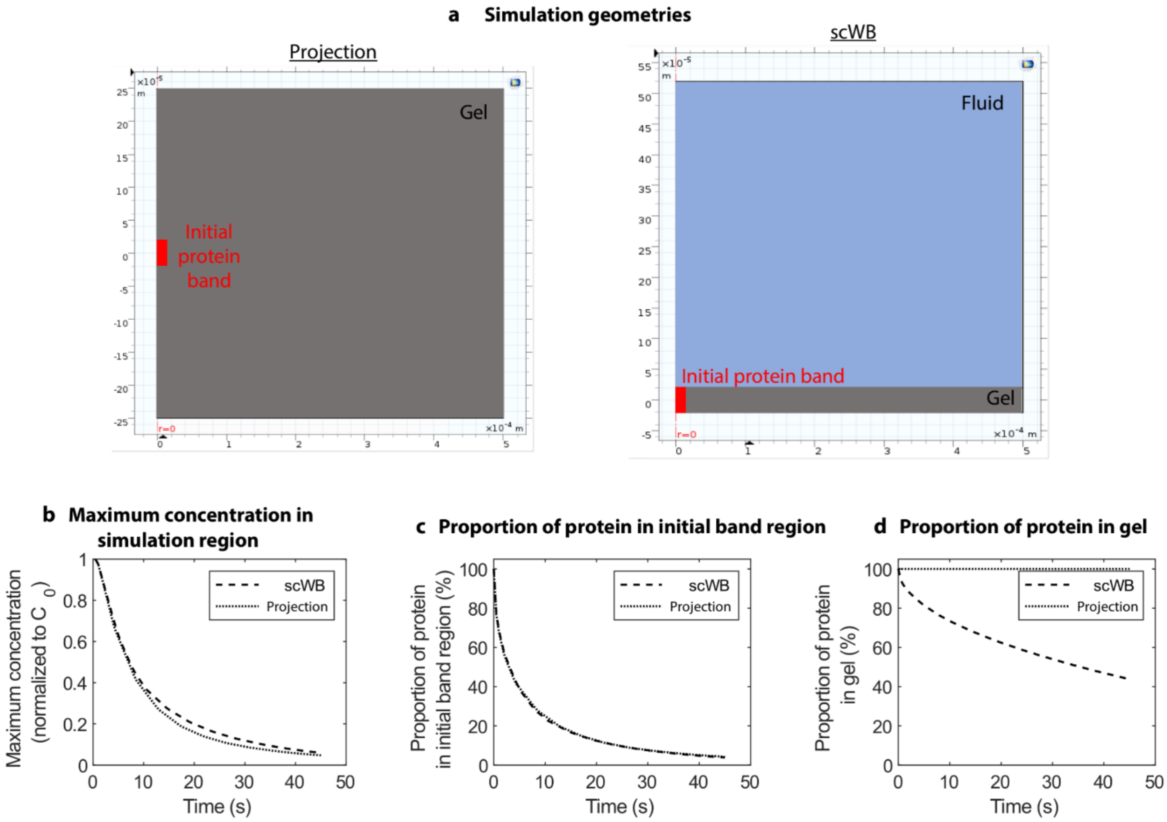


**Figure S2**. Comparison of simulated in-gel protein dilution during electrophoresis for standard single-cell western blotting and Z-direction electrophoresis. **(a)** 2D axisymmetric simulation geometries. **(b)** Comparison of maximum concentration in the simulation region, normalized to the initial concentration in the protein band ($C_0$). **(c)** Proportion of the protein in the initial simulation band region. **(d)** Proportion of protein retained in the gel.

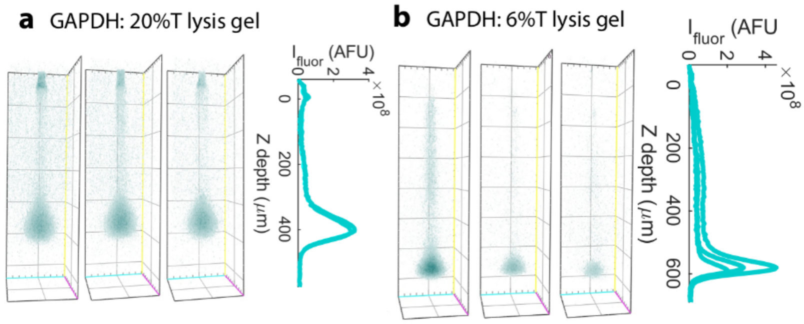


**Figure S3**. Representative 3D renderings (left) and summed fluorescence Z-intensity profiles (right) of GAPDH separations from BT474 breast tumour cells after lysis using **(a)** a 20%T lysis gel, and **(b)** a 6%T lysis gel, both using 2X RIPA + 8M urea lysis buffer and after 10s electrophoresis. By moving to 6%T lysis gels, we observed higher apparent GAPDH mobility (1.08±0.03x$10^{-4}$ $cm^2$/V·s using 6%T lysis gel, compared with 0.83±0.08x$10^{-4}$ $cm^2$/V·s using 20%T lysis gel, n=12-14 separation lanes) and potential reduction in dispersion of the protein band towards the microwell.

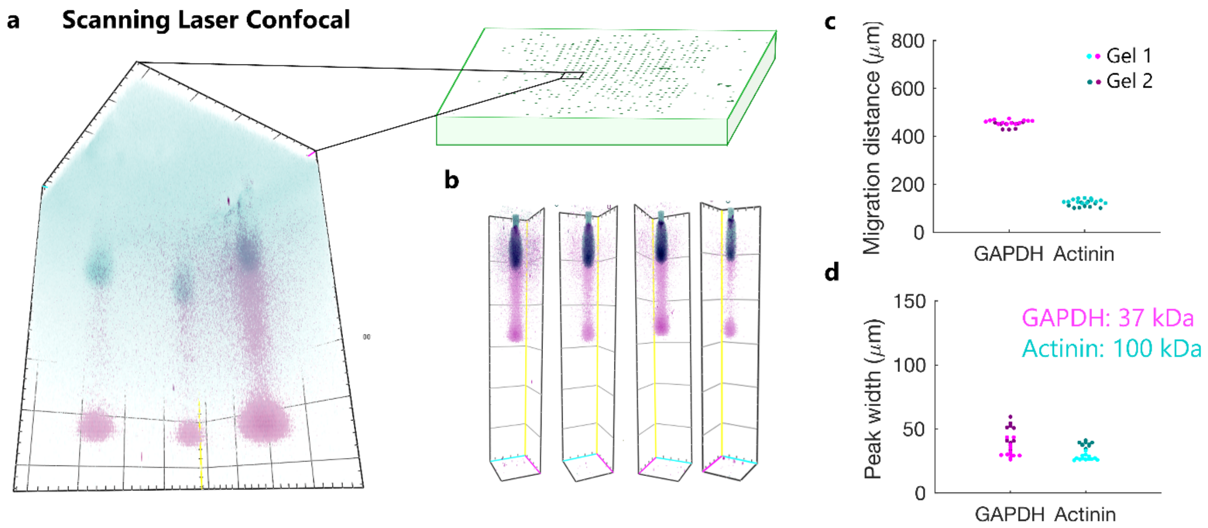


**Figure S4**. Comparison imaging by scanning laser confocal microscopy of the same projection electrophoresis separation gels analyzed in Figure 5. **(a)** a scanning laser confocal field of view compared to the size of the separation gel. **(b)** Representative individual separation lanes read out by scanning laser confical microscopy. **(c)** quantification of migration distance for GAPDH (37 kDa) and actinin (100 kDa). **(d)** quantification of Z-directional peak width for the separated bands of the same protein targets.

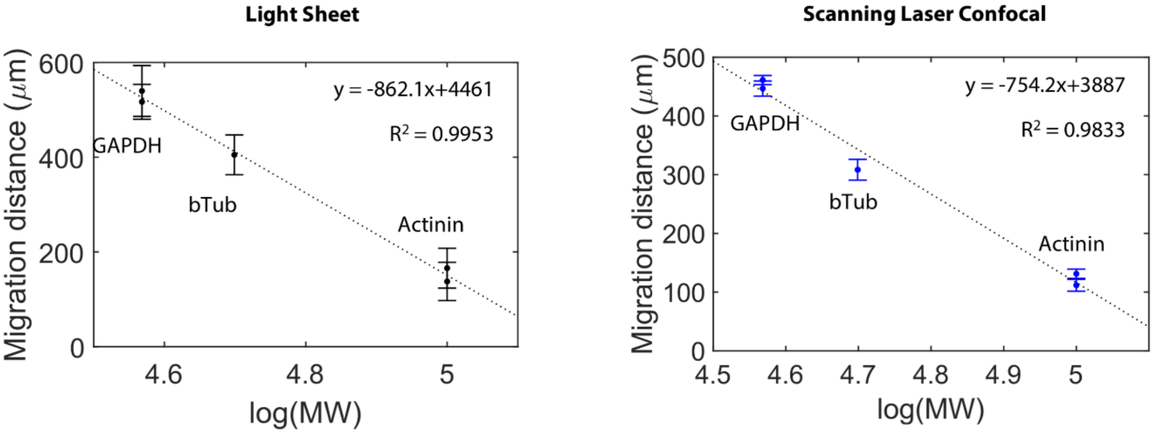


**Figure S5**. Quantification distance of migration distance vs. log(molecular weight) for three endogenous protein targets measured from single BT474 breast tumour cells using light sheet (left) and scanning laser confocal (right) microscopy readouts for the Projection Electrophoresis assay. Both readout methods show the expected log-linear relationship affirming size separation. Each point plots the mean and standard deviation of quantifiable separation lanes from a single separation gel (duplicate gels for GAPDH and actinin; a single gel for beta tubulin). For the light sheet analysis, 100-300 separation lines were quantified to yield each plotted point; for the scanning laser confocal analysis, 9-13 separation lanes were quantified to yield each plotted point.

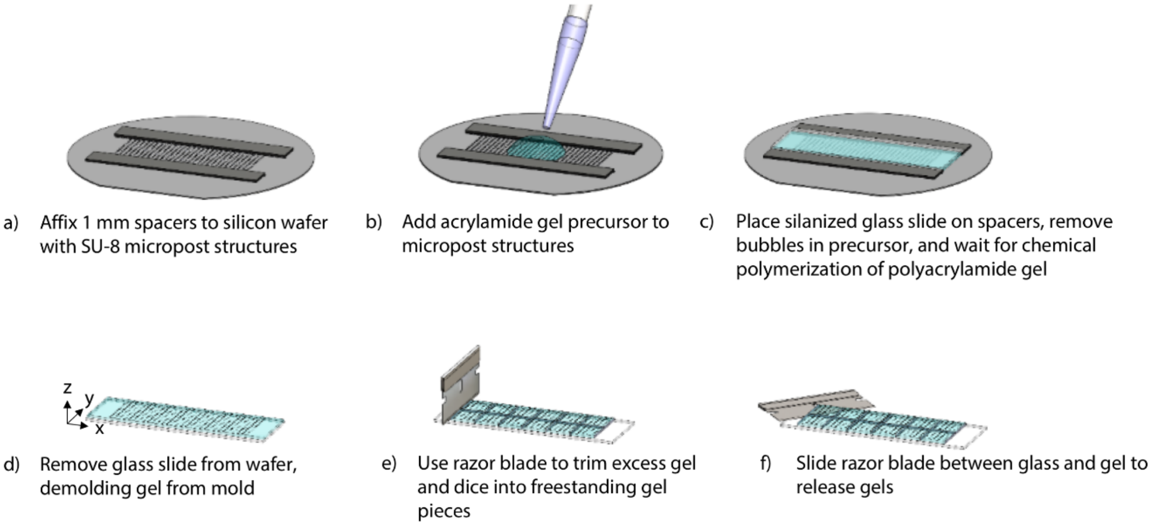


**Figure S6**. Substrate-free released gel fabrication process.

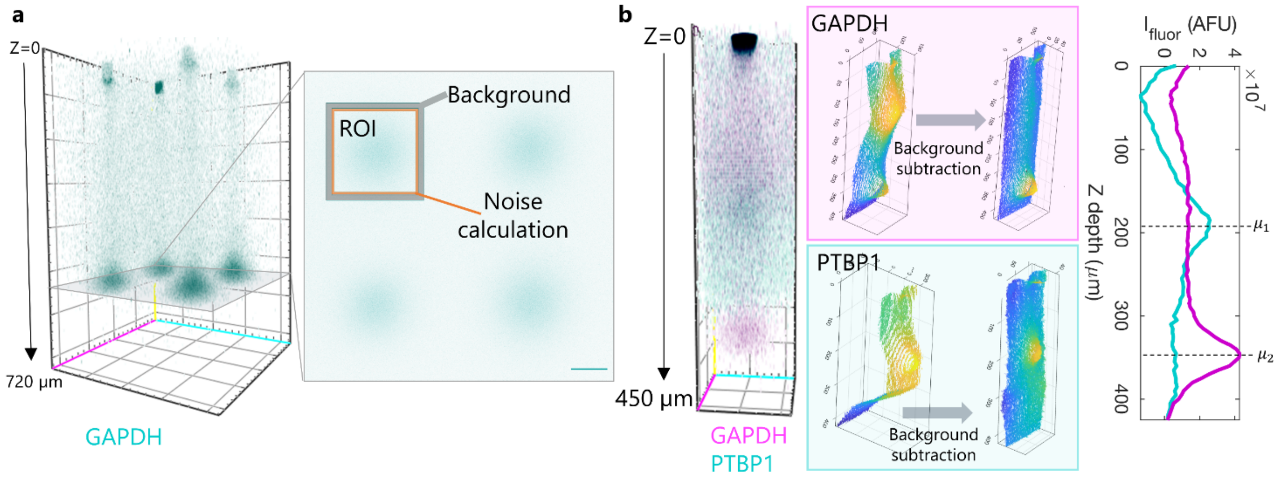


**Figure S7**. Data processing for three-dimensional projection electrophoresis datasets from immunoprobed separations from single BT474 cells. **(a)** 3D data is composed of stacks of X-Y slice images. Each slice image is processed to isolate the region of interest ('ROI') for each separation lane, as well as small adjacent surrounding regions for background subtraction ('Background') and to calculate the noise of the background-subtracted signal ('Noise calculation'). Scale bar represents 450 μm. **(b)** The ROI corresponding to each separation lane is also a 3D dataset (rendering, left), which can be collapsed into a 2D (X-Z) image by summing in Y (middle), or collapsed into a summed 1D intensity profile by summing all pixels in X and Y within the ROI region (right). Subtraction of the average background region intensity at each Z-depth from each pixel in the ROI (middle column) isolates the signal from the single-cell separation, clearly showing separated, immunoprobed protein peaks corresponding to GAPDH (top) and PTBP1 (bottom). Gaussian fitting to the background subtracted 1D intensity profiles yields migration distance, peak width, and signal-to-noise ratio information for each separation lane.

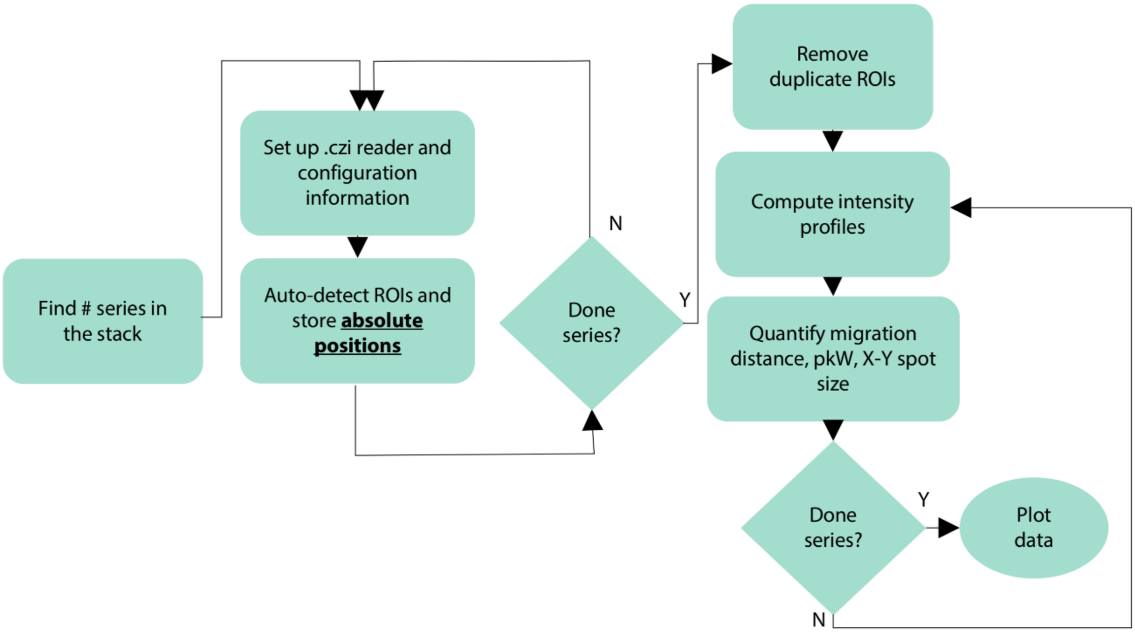


**Figure S8**. Schematic diagram of image analysis software for tiled light sheet images.

**Table S1.** Fabrication conditions for the various types of polyacrylamide gels used in this work.

| **Gel type (thickness)** | **Gel density** | **Stock acrylamide** | **Rhinohide** | **BPMA** | **Buffer** | **Initiator(s)** | **Polymerization time** | **Fabrication setup** |
|---|---|---|---|---|---|---|---|---|
| Purified protein: separation (1 mm) | 7%T or 10%T | 30% (37.5:1) stock; final concentration 7% or 10% w/v Sigma-Aldrich: A3699 | 10% v/v final from stock | 3 mM from 100 mM stock in DMSO | 10% (v/v) final concentration 10X tris-glycine; stored in modified RIPA | 0.08% APS (Sigma-Aldrich: A3678), 0.08% TEMED (Sigma-Aldrich: T9281) | 60 mins | Methacrylate functionalized glass slide and silanized silicon wafer mould |
| Purified protein: shield (500 µm) | 20%T | 30% (37.5:1) stock; final concentration 20% w/v | 10% v/v final from stock | none | 10% (v/v) final concentration 10X tris-glycine; stored in run buffer | 0.08% APS, 0.08% TEMED | 60 mins | Gel Slick® (Lonza: 50640) treated glass plate and glass slide |
| Well gels for single cell separations and in-well lysis tests (1 mm) | 7%T | 30% (37.5:1) stock; final concentration 7% w/v | 4.66% v/v final concentration from stock | 3 mM from 100 mM stock in DMSO | 10% (v/v) final concentration 10X tris-glycine; stored in PBS (lysis monitoring) or 1X tris-glycine (single cell separations) | 0.08% APS, 0.08% TEMED | 60 mins | Methacrylate functionalized glass slide and silanized silicon wafer mould |
| Lysis shield gels for in-well lysis tests (1 mm) | 20%T | 30% (37.5:1) stock; final concentration 20% w/v | 10% v/v final from stock | none | None; stored in modified RIPA | 0.08% APS, 0.08% TEMED | 60 mins | Gel Slick® treated glass plate and glass slide |
| Lysis shield gels for single-cell separations (1 mm) | 6%T or 20%T | 30% (37.5:1) stock; final concentration 6% or 20% w/v | 10% (20%T) or 4.66% (6%T) v/v final from stock | none | 10% (v/v) final concentration 10X tris-glycine; stored in 2X modified RIPA and transferred to 2X RIPA containing 8M urea for >10 minutes prior to separation. | 0.08% APS, 0.08% TEMED | 60 mins | Gel Slick® treated glass plate and glass slide |

**Table S2.** Projection electrophoresis buffers.

| **1X tris-glycine and 0.5% Triton X-100** | **1X tris-glycine** | **1X RIPA and 1X tris-glycine** | **2X RIPA and 2X tris-glycine** | **2X RIPA and 8M Urea** |
|---|---|---|---|---|
| 10X tris-glycine: 10% v/v Bio-Rad 1610734 | 10X tris-glycine: 10% v/v Bio-Rad 1610734 | 10X tris-glycine: 10% v/v Bio-Rad 1610734 | 10X tris-glycine: 20% v/v Bio-Rad 1610734 | Urea: 8M final (Sigma-Aldrich U5378) |
| Triton X-100: 0.5% v/v Sigma-Aldrich: X100 | MilliQ Water: 90% v/v | SDS: 0.5% w/v Sigma-Aldrich #L3771 | SDS: 1% w/v Sigma-Aldrich #L3771 | 10X tris-glycine: 20% v/v Bio-Rad 1610734 |
| MilliQ Water: 89.5% v/v | -- | Sodium Deoxycholate: 0.25% w/v Sigma-Aldrich #D6750 | Sodium Deoxycholate: 0.5% w/v Sigma-Aldrich #D6750 | SDS: 1% w/v Sigma-Aldrich #L3771 |
| -- | -- | Triton X-100: 0.1% v/v Sigma-Aldrich: X100 | Triton X-100: 0.2% v/v Sigma-Aldrich: X100 | Sodium Deoxycholate: 0.5% w/v Sigma-Aldrich #D6750 |
| -- | -- | MilliQ Water: 89.9% v/v | MilliQ Water: 79.8% v/v | Triton X-100: 0.2% v/v Sigma-Aldrich: X100 |
| -- | -- | -- | -- | MilliQ Water: 79.8% v/v |
